# Host kinase CSNK2 is a target for inhibition of pathogenic β-coronaviruses including SARS-CoV-2

**DOI:** 10.1101/2022.01.03.474779

**Authors:** Xuan Yang, Rebekah J. Dickmander, Armin Bayati, Sharon A. Taft-Benz, Jeffery L. Smith, Carrow I. Wells, Emily A. Madden, Jason W. Brown, Erik M. Lenarcic, Boyd L. Yount, Edcon Chang, Alison D. Axtman, Ralph S. Baric, Mark T. Heise, Peter S. McPherson, Nathaniel J. Moorman, Timothy M. Willson

## Abstract

Inhibition of the protein kinase CSNK2 with any of 30 specific and selective inhibitors representing different chemotypes, blocked replication of pathogenic human and murine β-coronaviruses. The potency of in-cell CSNK2A target engagement across the set of inhibitors correlated with antiviral activity and genetic knockdown confirmed the essential role of the CSNK2 holoenzyme in β-coronavirus replication. Spike protein uptake was blocked by CSNK2A inhibition, indicating that antiviral activity was due in part to a suppression of viral entry. CSNK2A inhibition may be a viable target for development of new broad spectrum anti-β-coronavirus drugs.

**GRAPHICAL ABSTRACT:** 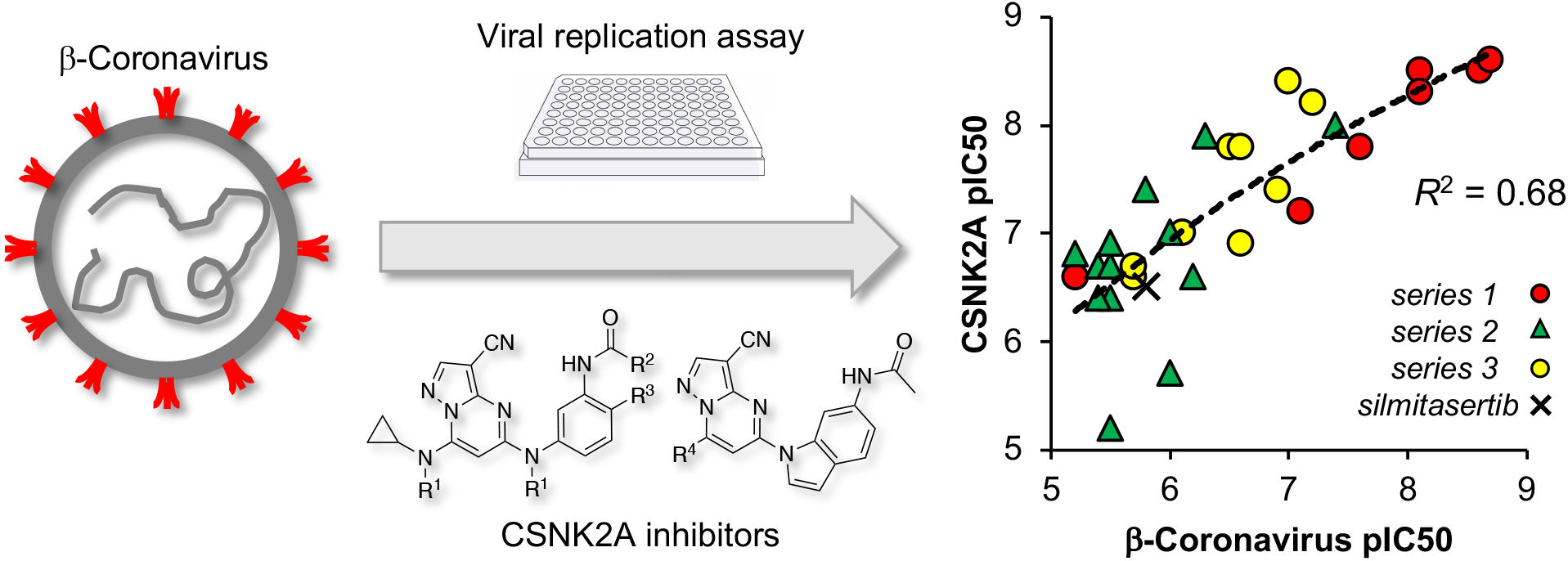

## INTRODUCTION

Coronaviruses (CoV) are genetically diverse positive-sense RNA viruses that circulate in animals and humans.^1^ α-CoV and β-CoV can infect mammals, while γ-CoV and δ-CoV are restricted to birds. Three highly pathogenic human β-CoV of zoonotic origin that cause severe lower respiratory tract infection have emerged in recent years: severe acute respiratory syndrome CoV (SARS-CoV), Middle East respiratory syndrome (MERS)-CoV, and SARS-CoV-2, the causative agent of the COVID-19 pandemic. Despite the rapid development of effective vaccines and direct-acting antivirals, the perpetual evolution of β-CoV, the inevitable emergence of drug resistance, and the potential for emergence of new zoonotic β-CoVs, have highlighted the need for effective broad-spectrum oral antiviral therapies to treat infections.^2^

CoVs are spherical enveloped viruses that are characterized by their crown-like surface projections, which are composed of trimers of the viral spike protein.^3^ The CoV spike protein binds receptors on the surface of target host cells to allow entry of the virus as the first step of infection. The CoV spike protein, which has adapted to target receptors of different hosts, determines the spectrum of infectivity of each virus. The spike proteins of SARS-CoV and SARS-CoV2 bind to human angiotensin converting enzyme 2 (ACE2) receptor,^4^ while MERS-CoV utilizes dipeptidyl peptidase 4^5^ and the spike protein of mouse hepatitis virus (MHV) binds to mouse carcinoembryonic antigen-related cell adhesion molecule 1 receptor.^6^ While SARS-CoV, SARS-CoV-2, and MERS-CoV are all biosafety level 3 pathogens, which makes work with these viruses more difficult, the restriction of MHV infectivity to mice, and its relation to other members of the β-CoV genus, makes it an excellent model system that can be studied with biosafety level 2 containment.^7^

Endocytosis of the receptor-bound virus is followed by release and uncoating of the CoV RNA genome.^1^ The virus encodes a replicase and an RNA-dependent RNA polymerase that transcribe the CoV mRNAs, which in turn are translated into the viral structural and nonstructural accessory proteins. Following assembly of new virions, composed of the viral genomic RNA and structural proteins, the virus is translocated in vesicles to the host cell membrane and released by nonlytic exocytosis. The virus coopts many host cell proteins through its life cycle to maintain efficient entry, replication, packaging, and exocytosis in addition to suppression of immune response pathways.^8^

Development of direct acting antiviral agents has often been hampered by the potential for viruses to overcome negative selective pressure to generate drug-resistant mutants.^9^ Host cell proteins that are utilized by the virus during replication or for suppression of the immune response are less likely to be circumvented by viral escape mutants.^10, 11^ Protein kinases are involved in almost all cell signaling processes and are often induced or suppressed by viruses during infection.^12^ Casein Kinase 2 (CSNK2) is a serine/threonine kinase typically found as a tetramer consisting of two catalytic subunits and two regulatory subunits, forming either a homotetramer or heterotetramer depending on the identity of the catalytic subunit (Figure 1A).^13^ A wide range of viruses have been shown to have proteins that are phosphorylated by CSNK2.^14^ It is unclear if all of these phosphorylation events are essential for virus replication or a manifestation of the broad range of CSNK2 substrate specificity. However, for human papillomaviruses, it appears that the phosphorylation of E1 protein by CSNK2 stabilizes ATP-dependent DNA helicase activity, which is key step in their viral replication.^15^ Recently a series of mass spectrometry proteomic and phosphoproteomic studies have mapped the interactions between β-CoV and host CSNK2 in infected cells.^16-18^ Both CSNK2A1 and CSNK2A2 were identified as participants in the SARS-CoV-2 interactome, specifically in a complex with the nucleocapsid protein.^16^ These observations were extended to SARS-CoV and MERS,^17^ suggesting that the interactions will be shared across other β-CoV members, such as MHV. Furthermore, phosphoproteomic profiling of cells following SARS-CoV-2 infection identified many CSNK2 substrates, which was consistent with upregulation of its kinase activity by the virus.^18^ These observations are indictive of β-CoV commandeering host cell CSNK2 to support its infectivity and replication and suggest that small molecule inhibitors may be promising antiviral compounds.

**Figure 1.**
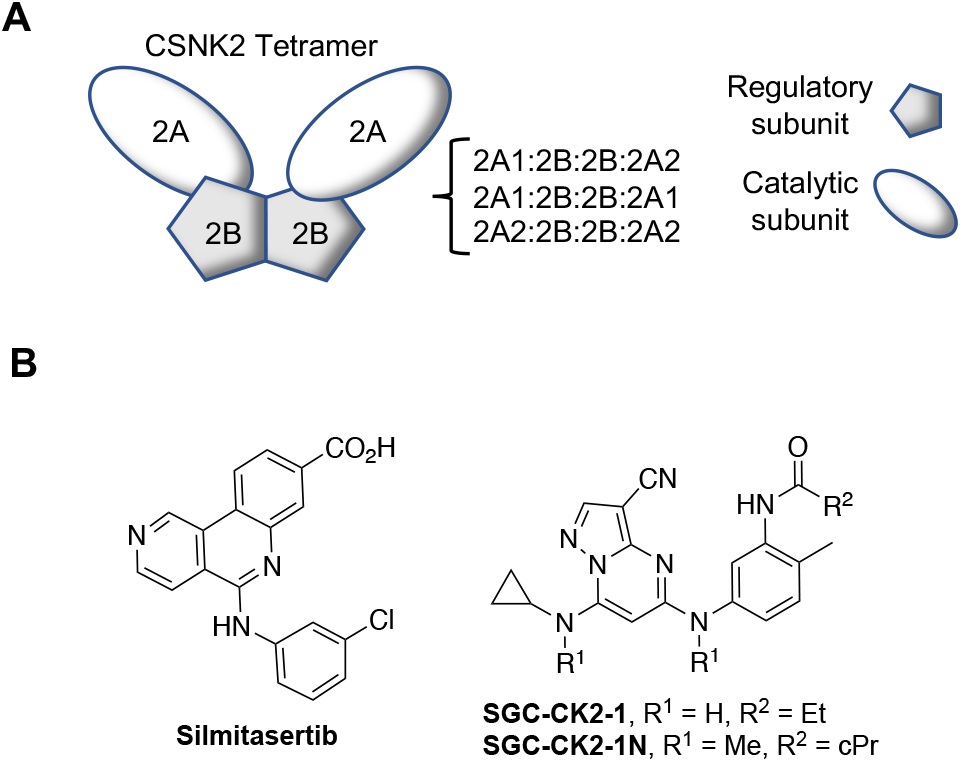
CSNK2 enzyme and inhibitors. (A) CSNK2 is a homo- or heterotetramer composed of two copies of the catalytic 2A-subunit (A1 or A2) and two copies of the regulatory 2B-subunit. (B) ATP-competitive CSNK2A inhibitors (silmitasetib and **SGC-CK2-1**) and negative control analog (**SGC-CK2-1N**).

Chemogenomics is a method of drug target validation that utilizes selective and highly annotated small molecule inhibitors to link perturbation of a cell phenotype to a specific molecular target.^19^ For protein kinases, a robust chemogenomic strategy requires the use of multiple small molecule inhibitor chemotypes combined with inactive analogs to control for potential pleiotropic kinase inhibition and other off-target activity.^20-22^ Well characterized ATP-competitive kinase inhibitors of CSNK2 have been developed, including silmitsertib and SGC-CK2-1 (Figure 1B). Using a chemogenomic approach employing multiple series of inhibitors we demonstrate that the potency of CSNK2A target engagement in cells tracks with the suppression of β-CoV replication. The critical role of CSNK2 in β-CoV replication was further confirmed by genetic knockdown of the individual catalytic and regulatory subunits. Finally, by studying the effect of CSNK2A inhibition on SARS-CoV-2 spike protein uptake we provide evidence that antiviral activity is manifest in part by inhibition of viral entry into cells.

## RESULTS AND DISCUSSION

### Development of a β-coronavirus reporter assay

Mouse hepatitis virus (MHV) is a member of the β-CoV genus that has been widely used as a model to study the virulence of SARS-CoV and SARS-CoV-2.^7^ To develop a reporter virus to study the effect of compounds on β-CoV replication, the MHV-A59 G plasmid was engineered to replace most of the coding sequence for orf4a and orf4b with nanoluciferase (nLuc).^23^ The resulting virus, MHV-nLuc, replicated to high titer and efficiently expressed nLuc.

To determine the optimal titer and time point to analyze viral replication, mouse derived-from-brain-tumor (DBT) cells were inoculated with a range of multiplicities of infection (MOI) from 0.016 to 10 with MHV-nLuc and luciferase activity measured in cell lysates at multiple time points up to 24 h post infection (Figure 2A). The results indicated that inoculation of DBT cells by MHV-nLuc with an MOI of 0.1 and luciferase measurement at 10 h post infection were the optimal assay conditions, as viral replication was in the linear range and bioluminescence was within the dynamic range of the luminometer.

**Figure 2.**
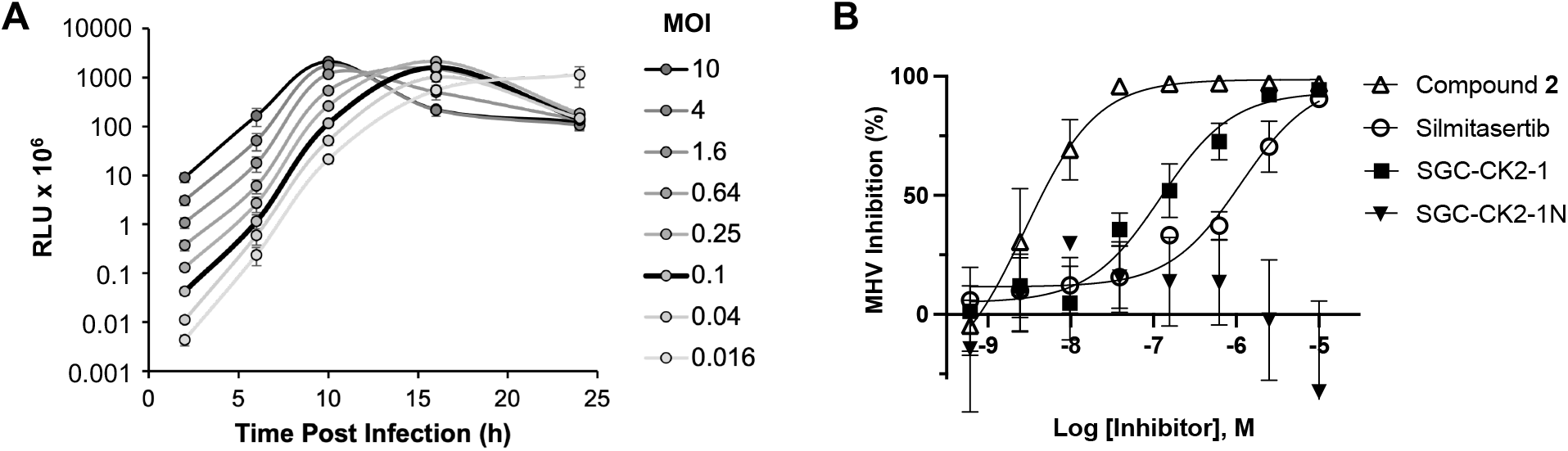
β-CoV replication assay. (A) Optimization of MHV-nLuc assay. (B) Effect of CSNK2A inhibitors on replication of MHV-nLuc in DBT cells, n = 3 ± SE. No curve was fit to the **SGC-CK2-1N** data.

### CSNK2A inhibitors block β-CoV replication

Silmitasertib (Figure 1B) is a modestly selective ATP-competitive CSNK2 inhibitor with a live cell CSNK2A1 target engagement of pIC_50_ = 6.5 as measured by nanoBRET assay.^24, 25^ Silmitasertib was previously reported as demonstrating antiviral activity in African green monkey kidney epithelial Vero cells infected with SARS-CoV-2 (pIC_50_ = 5.6).^18^ However, it was less potent at inhibiting SARS-CoV-2 infection of human lung epithelial A549-ACE2 cells (pIC_50_ <5). Silmitasertib demonstrated cell toxicity at micromolar doses in both cell lines,^18^ which complicated analysis of its anti-SARS-CoV-2 activity. When tested in our optimized MHV-nLuc assay in DBT cells silmitasertib inhibited viral replication with pIC_50_ = 6.2 and with no effect on cell viability (Figure 2B and SI Figure 1). These data demonstrated for the first time that the anti-β-CoV activity of silmitasertib could be uncoupled from its effect on cell viability.

To provide additional evidence that host cell CSNK2 was required for coronavirus replication, we tested a second series of ATP-competitive CSNK2A inhibitors from the pyrazolo[1,5-*a*]pyrimidine chemotype which is structurally and physiochemically distinct from silmitasertib (Table 1). We previously reported the identification of a series of 3-cyano-7-cyclopropylamino-pyrazolo[1,5-*a*]pyrimidines **1**–**7** as potent, selective, cell active inhibitors with pIC_50_ = 6.6–8.9 in CSNK2A1/CSNK2A2 nanoBRET assays.^24^ When tested in the MHV-nLuc assay, pyrazolo[1,5-*a*]pyrimidines **1**–**6** demonstrated potent inhibition of viral replication. Pyrazolo[1,5-*a*]pyrimidines **1**–**4** demonstrated IC_50_ values <10 nM with no effect on viability of DBT cells at concentrations up to 10 μM (SI Figure 1). The *N*-benzyl pyrazolo[1,5-*a*]pyrimidine **7** was the only analog with an IC_50_ above 1 μM. The lower potency of **7** in the MHV-nLuc assay was consistent with its weaker activity in the CSNK2A1/CSNK2A2 nanoBRET assays.

**Table 1.** Structure-activity of the *N*-(3-aminophenyl)acetamide series of 3-cyano-7-cyclopropylamino-pyrazolo[1,5-*a*]pyrimidines **1**–**7**.

| Compound | R | CSNK2A1<br>nanoBRET<br>($pIC_{50}$)<sup>a</sup> | CSNK2A2<br>nanoBRET<br>($pIC_{50}$)<sup>a</sup> | MHV<br>replication<br>($pIC_{50}$)<sup>b</sup> |
| --- | --- | --- | --- | --- |
| **silmitasertib** | — | 6.5 | 7.3 | 6.2 |
| **1** |  | 8.5 | 8.7 | 8.1 |
| **2** |  | 8.5 | 8.6 | 8.6 |
| **3** |  | 8.3 | 8.4 | 8.1 |
| **4** |  | 8.6 | 8.9 | 8.7 |
| **5** |  | 7.8 | 8.1 | 7.6 |
| **6** |  | 7.2 | 7.8 | 7.1 |
| **7** |  | 6.6 | 6.9 | 5.2 |

<sup>a</sup> In cell target engagement of CSNK2A-nLuc in HEK293 cells. Data from reference 24. <sup>b</sup> Inhibition of MHV-nLuc replication in DBT cells. Values are the mean of three assays with range $\pm 15\%$ .

The kinome-wide selectivity of the 3-cyano-7-cyclopropylamino-pyrazolo[1,5-*a*]pyrimidine inhibitors is controlled in part by the *para*- and *meta*-aniline substituents.^24^ **SGC-CK2-1** (Figure 1B), which contains *para*-methyl and *meta*-propionamide aniline substituents, is the most selective of all known ATP-competitive small molecule CSNK2A inhibitors (3 kinases inhibited at 1 μM)^24^ and has been characterized as a high-quality chemical probe by the Structural Genomics Consortium.^22^ A close structural analog, **SGC-CK2-1N** (Figure 1B), that lacks CSNK2A activity at concentrations up to 10 μM has been designated as a negative control compound. When tested in the MHV-nLuc assay **SGC-CK2-1** inhibited viral replication with pIC_50_ = 6.9 while negative control **SGC-CK2-1N** was inactive up to a concentration of 10 μM (Figure 2B). This result provided a third line of chemogenomic evidence that inhibition of host cell CSNK2 impeded replication of a β-CoV.

### Relationship between CSNK2A inhibitor potency and anti-β-CoV activity

To generate additional evidence that CSNK2 was required for β-CoV replication, two additional series of inhibitors based on the pyrazolo[1,5-*a*]pyrimidine chemotype were synthesized to strengthen the structure-activity relationship between kinase inhibition and viral replication (Scheme 1). The 3-cyano-7-cyclopropylamino-pyrazolo[1,5-*a*]pyrimidines, in particular, have demonstrated high cellular potency as CSNK2A inhibitors combined with good kinome-wide selectivity.^24^ A series of analogs (**8**–**15**, Table 2) where the aniline *para*-methyl group of the chemical probe **SGC-CK2-1** was replaced by a basic sidechain were synthesized by palladium catalyzed cross-coupling of di-substituted aniline intermediates (**i**) and pyrazolo[1,5-*a*]pyrimidine building block (**ii**) (Scheme 1). The *para*-substituent on the aniline forces the propionamide to adopt an otherwise energetically disfavored cisoid configuration in the enzyme active site that contributes to improved CSNK2A selectivity.^24^ Although the ATP-binding sites of CSNK2A1 and CSNK2A2 have high sequence identity (SI Figure 2), we opted to screen the new analogs **8**–**15** for cellular target engagement on both isozymes using nanoBRET assays. MHV replication tracked with nanoBRET activity, with the most potent dual CSNK2A1/CSNK2A2 inhibitors **8** and **9** showing Vthe strongest MHV inhibition and the least effective inhibitors **12** and **14** showing the weakest inhibition of viral replication. Notably analogs **10** and **15** showed modest selectivity for CSNK2A1 over CSNK2A2 (6–8 fold), but this did not translate into improved potency for MHV inhibition.

**Table 2.**
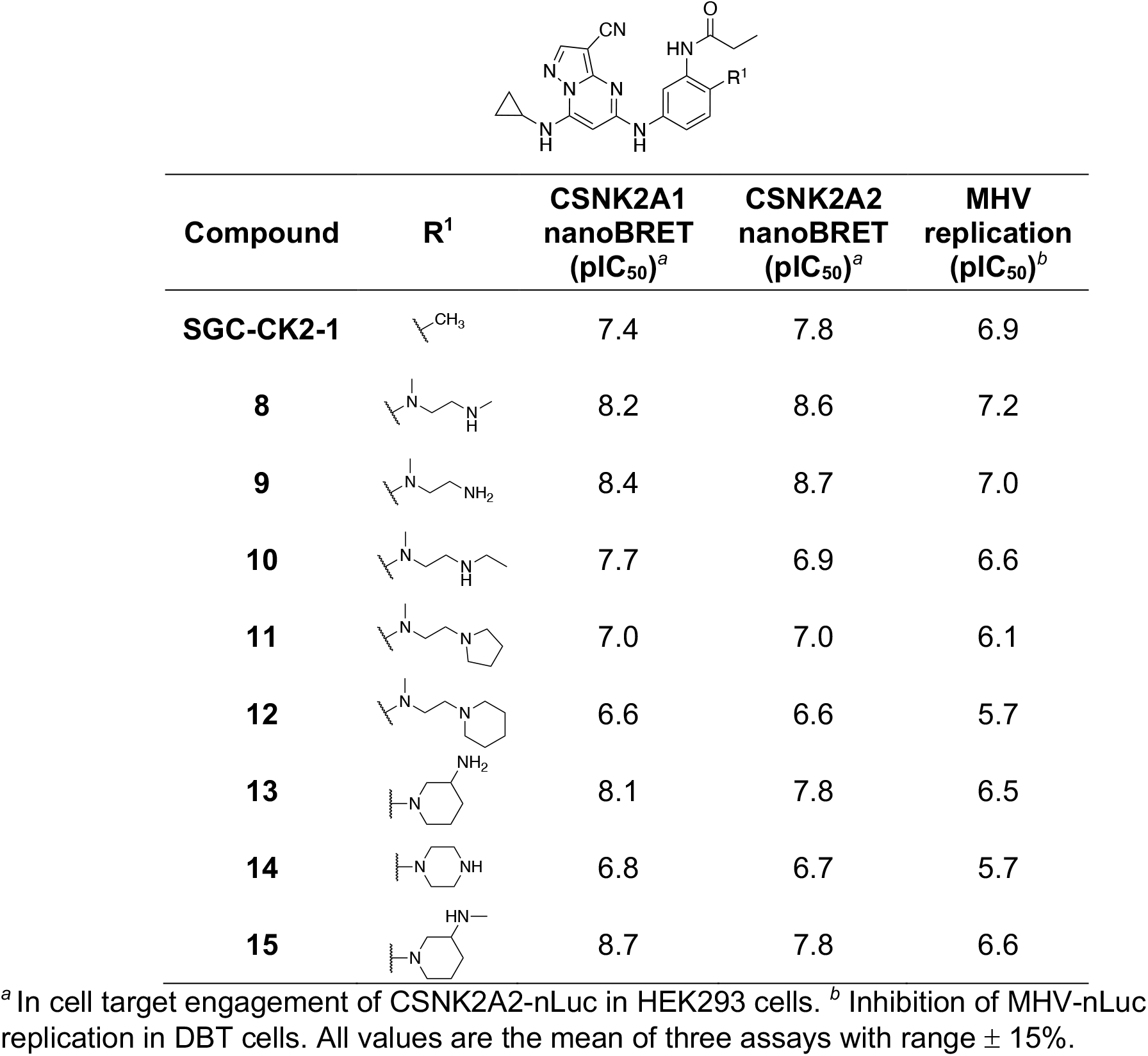
Structure-activity of the *N*-(3-aminophenyl)propionamide series of 3-cyano-7-cyclopropylamino-pyrazolo[1,5-*a*]pyrimidines **8-15**.

| Compound | R<sup>1</sup> | CSNK2A1<br>nanoBRET<br>(pIC<sub>50</sub>)<sup>a</sup> | CSNK2A2<br>nanoBRET<br>(pIC<sub>50</sub>)<sup>a</sup> | MHV<br>replication<br>(pIC<sub>50</sub>)<sup>b</sup> |
| --- | --- | --- | --- | --- |
| **SGC-CK2-1** |  | 7.4 | 7.8 | 6.9 |
| **8** |  | 8.2 | 8.6 | 7.2 |
| **9** |  | 8.4 | 8.7 | 7.0 |
| **10** |  | 7.7 | 6.9 | 6.6 |
| **11** |  | 7.0 | 7.0 | 6.1 |
| **12** |  | 6.6 | 6.6 | 5.7 |
| **13** |  | 8.1 | 7.8 | 6.5 |
| **14** |  | 6.8 | 6.7 | 5.7 |
| **15** |  | 8.7 | 7.8 | 6.6 |

<sup>a</sup> In cell target engagement of CSNK2A2-nLuc in HEK293 cells. <sup>b</sup> Inhibition of MHV-nLuc replication in DBT cells. All values are the mean of three assays with range ± 15%.

**Scheme 1.**
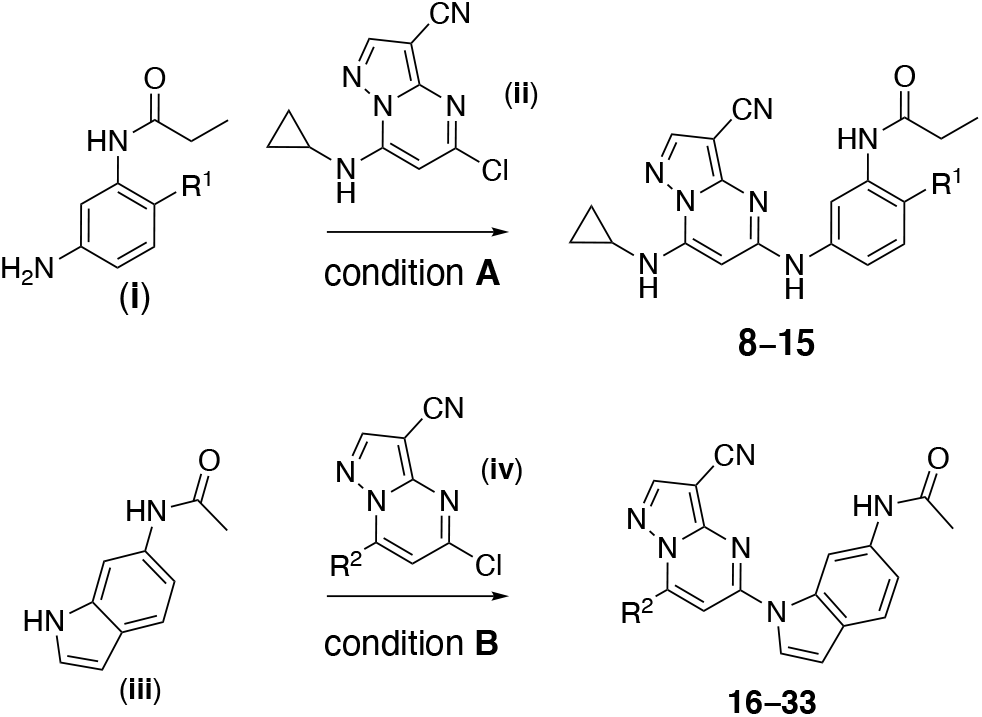
Synthesis of 3-Cyano-pyrazolo[1,5-*a*]pyrimidines **8**–**33**^*a*^. ^*a*^ Reagents and conditions: (A) BINAP, Pd(OAc)_2_, *t*-BuOLi, 1,4-dioxane, microwave irradiation. R^1^ defined in Table 2. (B) Xantphos, Pd(OAc)_2_, Cs_2_CO_3_, 1,4-dioxane, microwave irradiation, 130 °C 130 °C. R^2^ defined in Table 3.

The third series of a potent CSNK2A inhibitors was based on pyrazolo[1,5-*a*]pyrimidine **16** which contained a 6-(acetamino)indole as its 5-substituent.^24^ Indole **16** also demonstrated potent anti-β-CoV activity with pIC_50_ = 7.4 in the MHV-nLuc assay (Table 3). Guided by the knowledge that the cyclopropylamine sits in the region of the kinase that is adjacent to the solvent accessible channel, we explored a range of alternative 7-heterosubstituted analogs to define the structure-activity for CSNK2A inhibition and anti-β-CoV activity (Table 3). Synthesis of the analogs **16**–**33** was achieved by palladium catalyzed cross-coupling of 6-acetaminoindole (**iii**) and 7-substituted choloro-pyrazolo[1,5-*a*]pyrimidines (**iv**) (Scheme 1). In general, large modifications to the 7-cyclopropylamino group were found to be deleterious to in cell CSNK2A1/CSNK2A2 target engagement, but smaller modifications retained activity on the enzyme (Table 3). Importantly, as was seen with the aniline-substituted pyrazolo[1,5-*a*]pyrimidines, anti-β-CoV activity of the 6-(acetamino)indoles tracked with their cellular potency on CSNK2A. Only the 7-cyclobutylamino analog **17** demonstrated a pIC_50_ >6.0 in the MHV-nLuc assay. However, several of the analogs that had modest potency in the CSNK2A1/CSNK2A2 nanoBRET assays demonstrated IC_50_ values in the 1–5 μM range in the MHV-nLuc assay. Importantly, the analogs **21** and **32** that were inactive at 10 μM on CSNK2A were unable to block β-CoV replication. As was seen in before, analogs with modest CSNK2A1 selectivity (e.g. **25** and **28**) did not show improved anti-viral potency. Thus, even though the compounds in the 6-(acetamino)indole series were generally less active as dual CSNK2A1/CSNK2A2 inhibitors, their potency tracked with antiviral activity.

**Table 3.** Structure-activity of the *N*-(1*H*-indol-6-yl)acetamide series of 3-cyanopyrazolo[1,5-*a*]pyrimidines **16**–**33**.

| Compound | R<sup>2</sup> | CSNK2A1<br>nanoBRET<br>(pIC<sub>50</sub>)<sup>a</sup> | CSNK2A2<br>nanoBRET<br>(pIC<sub>50</sub>)<sup>a</sup> | MHV<br>replication<br>(pIC<sub>50</sub>)<sup>b</sup> |
| --- | --- | --- | --- | --- |
| **16** |  | 8.0 | 8.4 | 7.4 |
| **17** |  | 7.9 | 8.0 | 6.3 |
| **18** |  | 7.4 | 7.0 | 6.0 |
| **19** |  | 6.8 | 7.1 | 5.2 |
| **20** |  | 6.3 | 5.8 | i.a. |
| **21** |  | i.a. | i.a. | i.a. |
| **22** |  | 5.1 | i.a. | i.a. |
| **23** |  | 7.4 | 7.6 | 5.8 |
| **24** |  | 7.0 | 6.7 | 5.4 |
| **25** |  | 7.5 | 6.6 | 6.2 |
| **26** |  | 7.0 | 5.7 | 6.0 |
| **27** |  | 6.4 | 6.7 | 5.5 |
| **28** |  | 7.2 | 6.4 | 5.4 |
| **29** |  | 6.8 | 6.7 | 5.5 |
| **30** |  | 6.9 | 7.1 | 5.5 |
| **31** |  | 5.4 | 5.9 | i.a. |
| **32** |  | i.a. | i.a. | i.a. |
| **33** |  | 5.2 | 5.2 | 5.5 |

<sup>a</sup> In cell target engagement of CSNK2A2-nLuc in HEK293 cells. <sup>b</sup> Inhibition of MHV-nLuc replication in DBT cells. All values are the mean of three assays with range $\pm 15\%$ . i.a. inactive.

### Relationship between CSNK2A potency and β-CoV replication

Over the three series of 3-cyano-pyrazolo[1,5-*a*]pyrimidines (Tables 1–3) with a wide range of 5- and 7-substituents anti-β-CoV activity shadowed their potency in the live cell CSNK2A1/CSNK2A2 target engagement assays (Figure 3). Within each series the most potent kinase inhibitors were the most potent in the antiviral assay and the least active CSNK2A inhibitors unable to block viral replication. The relationship was maintained over a nearly 4 log range in activity with an *R*^2^ = 0.68 when the lower value for inhibition of CSNK2A1 or CSNK2A2 was compared to anti-viral potency (Figure 3). The relationship was also maintained with *R*^2^ > 0.6 when either CSNK2A1 or CSNK2A2 alone were used in the analysis (SI Figure 2). However, the improved correlation obtained when using target engagement data from both catalytic isoforms suggests that the heterotetramer form of the holoenzyme (Figure 1A) is the active complex in cells and that dual CSNK2A1/CSNK2A2 inhibition translates to improved anti-viral potency. The modest potency of silmitasertib, which belongs to a different chemotype of CSNK2A inhibitors, was also consistent with the relationship between CSNK2A and anti-β-CoV activity. The *N*-(3-aminophenyl) acetamide series (Table 1) contained the most potent inhibitors of CSNK2A and MHV-NLuc.

**Figure 3.**
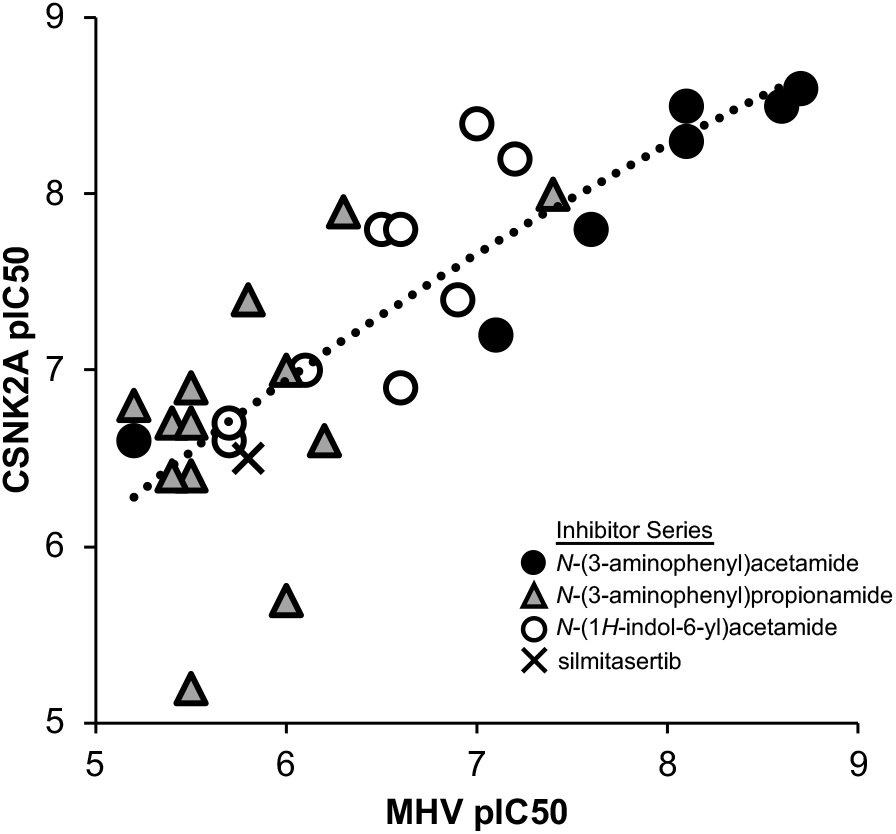
Correlation of potency for CSNK2A target engagement with inhibition of β-CoV replication across three subseries of pyrazolo[1,5-*a*]pyrimidines and silmitasertib. The analysis uses the lower value of CSNK2A1 or CSNK2A2 pIC_50_ from Tables 1–3 for each active analog. Inactive analogs **SGC-CK2-1N, 20, 21, 22, 31**, and **32** were not included in the analysis. The dashed line represents the logarithmic trendline with *R*^2^ = 0.68.

Several analogs in the *N*-(3-aminophenyl)propionamide series (Table 2) maintained potent CSNK2A activity with many analogs showing activity <1 μM, but no single analog as potent in the β-CoV replication assay as members of the *N*-acetamide series. Thus, while paired analogs had equivalent activity in the CSNK2A nanoBRET assays, the propionamide series was generally less potent in the antiviral assay (see **1** vs **SGC-CK2-1, 2** vs **8**, and **3** vs **9**). These nuances in the structure-activity relationship are unlikely to be due to species differences between the human kinase and the murine viral assay, since both human and mouse CSNK2A1 are identical in the kinase domain and CSNK2A2 differs by only a single amino acid E253D at the base of the C-lobe at >30 Å from the ATP-binding pocket (SI Figure 3). Physiochemical properties can also contribute to non-enzymatic viral inhibition mechanisms such as phospholipidosis.^26^ This non-specific activity is unlikely to confound our results due to the strong correlation between CSNK2A activity and MHV inhibition across a wide dose range and the nanomolar potency of many of the CSNK2A inhibitors. However, we cannot rule out some potential non-specific mechanisms with the two weakest CSNK2A inhibitors (pIC50 <6) that lie the furthest from the trendline (Figure 3).

### Targeted knock down of CSNK2 blocks β-CoV replication

CSNK2 is a serine/threonine kinase that is expressed endogenously as a tetramer of two catalytic subunits and two regulatory subunits, forming either a homotetramer or heterotetramer depending on the identity of the catalytic subunit (Figure 1A).^13^ Transcripts for each of the three subunits (CSNK2A1, CSNK2A2, CSNK2B) were detected in uninfected DBT cells by qRT-PCR. To study the effect of MHV infection, DBT cells were inoculated at an MOI of 0.1 and the transcript abundance of each CSNK2 subunit was determined through the time course of infection. By qRT-PCR, MHV infection did not change the abundance of any CSNK2 subunit transcripts in DBT cells over 12 hours (Figure 4A).

**Figure 4.**
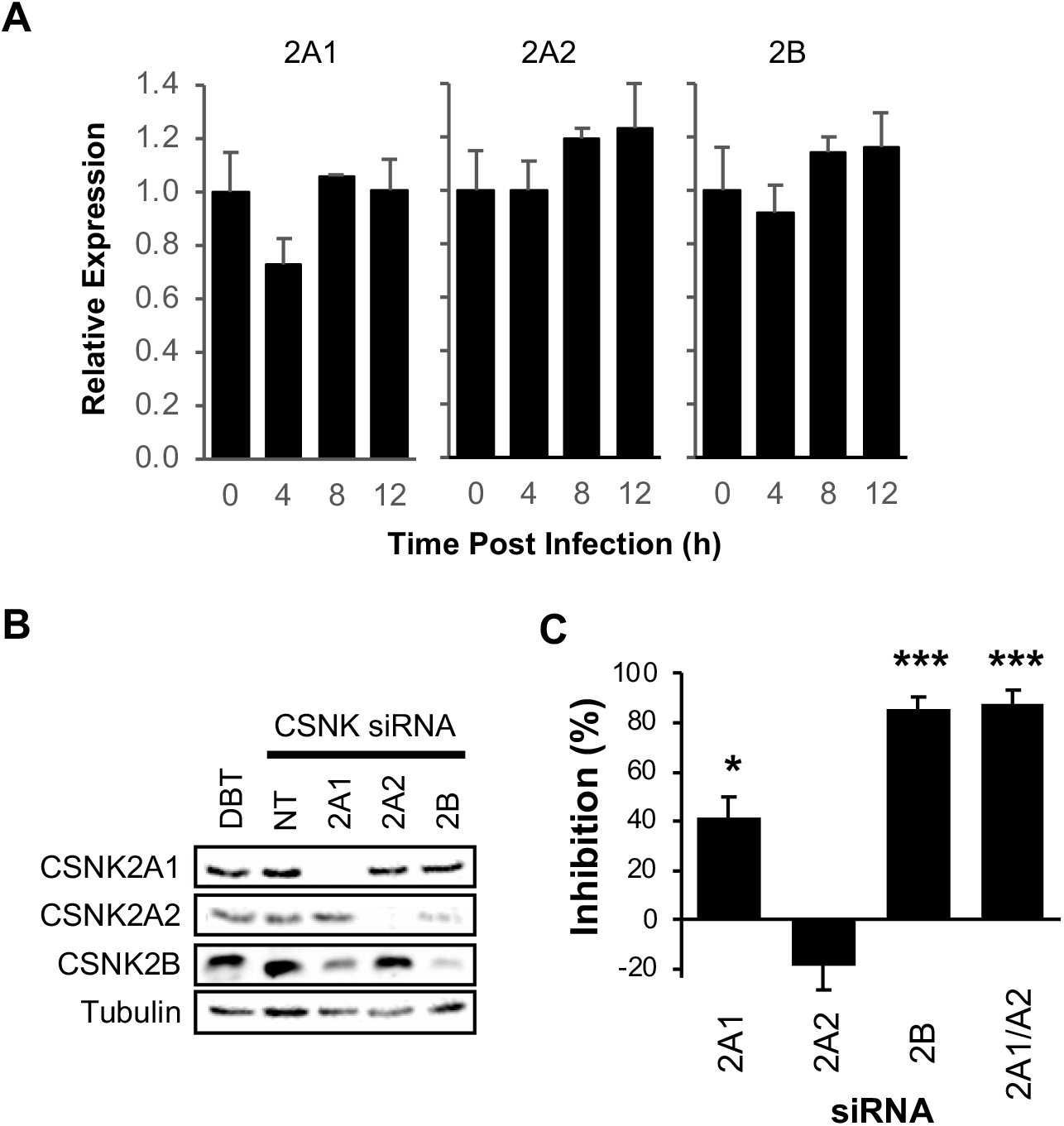
CSNK2 target validation. (A) Relative expression after MHV infection of the CSNK2 subunit mRNAs in DBT cells by qRT-PCR. (B) Expression of CSNK2 subunits by Western blot after siRNA targeting. DBT, untreated cells. NT, non-targeting control siRNA. **C**. Inhibition of MHV replication by siRNA knockdown of CSNK2 subunits. * *p* < 0.05, *** *p* < 0.001.

To further validate the role of CSNK2 in supporting β-CoV replication, targeted knockdown of the individual subunits of the enzyme was performed with siRNA specific to CSNK2A1, CSNK2A2, or CSNK2B respectively. Effective knockdown of each subunit was confirmed by Western blot using a well characterized antibody (Figure 4B).^24^ Following knockdown, the DBT cells were infected with MHV-nLuc at an MOI of 0.1 to determine the role of each subunit on viral replication. Knockdown of CSNK2A1 inhibited MHV replication by 40% compared to a non-targeting control siRNA, whereas CSNK2A2 knockdown did not inhibit MHV replication.

Knockdown of CSNK2B inhibited MHV replication by 85% compared to the control siRNA (Figure 4C). These results support the model (Figure 1A) in which a functional CSNK2 tetramer can be assembled using two copies of either CSNK2A1 or CSNK2A2 but must always contain two CSNK2B subunits. Thus, while the two catalytic subunits can be any mix of CSNK2A1 and CSNK2A2 subunits, the absence of CSNK2B yields a non-functional enzyme and loss of MHV replication in the DBT cells. To confirm our interpretation of the results, both CSNKA1 and CSNKA2 were depleted simultaneously. Dual knockdown of CSNK2A1 and CSNK2A2 inhibited MHV replication by 90% compared to the control siRNA (Figure 4B), further supporting the critical role of both isoforms of the catalytic unit during β-CoV replication. Evidence that the 2A1:2B:2B:2A2 heterotetramer is likely to be the primary form of the CSNK2 holoenzyme in cells was also provided by the chemogenomic analysis, which showed a stronger correlation with inhibition of β-CoV replication using target engagement data from both the CSNK2A1 and CSNK2A2 catalytic units (Figure 3 and SI Figure 2) and the observation that inhibitors with modest selectivity for CSNK2A1 over CSNK2A2 did not show improved anti-viral potency.

### CSNK2 inhibition blocks SARS-CoV-2 replication

To extend these findings to the clinically relevant SARS-CoV-2 that is the cause of the COVID-19 pandemic, we studied the effect of CSNK2 inhibition on virus replication in both continuous cell lines and in primary human cells. The severe contagion risk of SARS-CoV-2 mandates the use of high containment biosafety laboratory 3 containment for these assays, which restricted the study to the potent pyrazolo[1,5-*a*]pyrimidine CSNK2A inhibitor **2** (Figure 2B and Table 1). When tested in A549-ACE2 cells infected with a previously described SARS-CoV-2 virus expressing n-Luc,^27^ a dose dependent inhibition of viral replication was observed with 100% inhibition at concentrations above 700 nM (Figure 5A). However, viability of the A549-ACE2 cells was also impacted at higher doses. It is unclear why ATP-competitive CSNK2A inhibitors in two different chemotypes negatively impact viability of A549-ACE2 cells at low micromolar concentrations. However, it is known that silmersertib shows collateral inhibition of the CLK family of kinases that contribute to its antiproliferative activity, while the pyrazolo[1,5-*a*]pyrimidines show cross-activity on the DAPK family of kinases at micromolar concentrations.^24^ Since toxicity may be a manifestation of the transformed A549-ACE2 cell line and to provide additional evidence that CSNK2 inhibition negatively impacts SARS-CoV-2 replication, we measured the effect of CSNK2 inhibitors on SARS-CoV-2 replication in primary human airway epithelial cells (HAE) grown in culture on an air-liquid interface. These primary lung cells model the architecture and cellular complexity of the conducting airway and are readily infected by zoonotic CoV, including SARS-CoV-2.^27-29^ At a dose of 10 μM, CSNK2A inhibitor **2** caused a 1.5 to 2-log reduction in the level of SARS-CoV-2 in HAE derived from three different donors after 18 hours without affecting cell viability (Figure 5B). The efficacy of **2** was equivalent to remdesivir dosed at a 2.5 μM concentration. A dose response assay in the HAE cells indicated that **2** had an IC_50_ in the 200–300 nM range for inhibition of SARS-CoV-2 replication without affecting cell viability at doses up to 10 μM (Figure 5C). Combined with the results from the MHV-nLuc assay, these data provide strong evidence of the efficacy of host cell CSNK2A inhibitors in preventing replication of β-CoV, including SARS-CoV-2.

**Figure 5.**
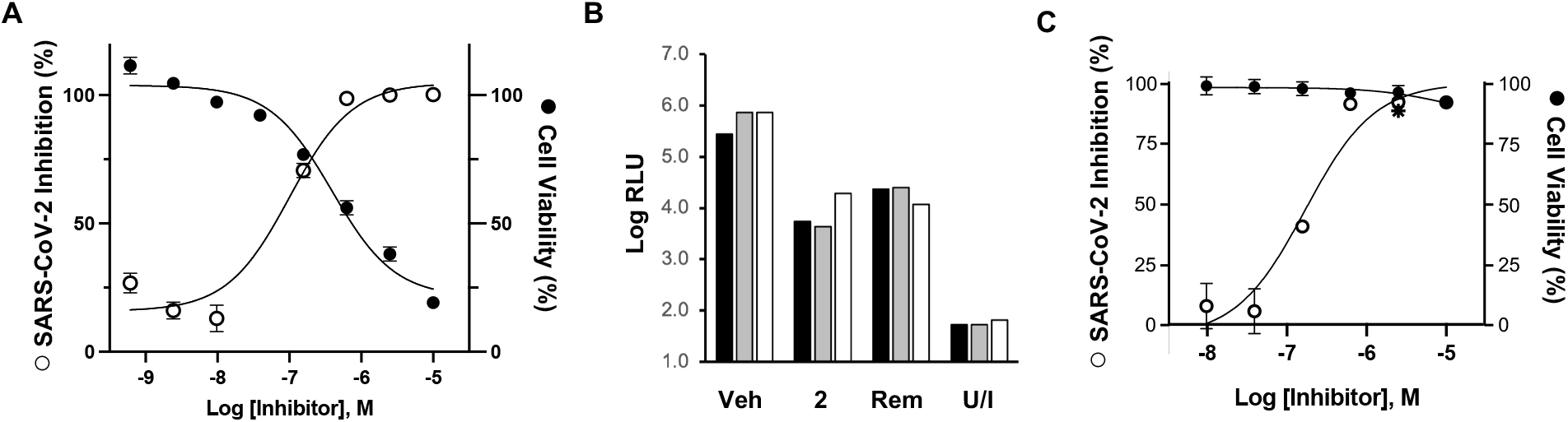
SARS-CoV-2 replication assays. (A) A549-ACE2 cells infected with SARS-CoV-2-nLuc. Effect of CSNK2A inhibitor **2** on inhibition of viral replication (open circles). Cell viability determined by LDH assay (closed circles). Assay performed in triplicate with data ranges shown by error bars. (B) Primary HAE cells from three donors (black, grey, while bars) infected with SARS-CoV-2-nLuc or unifected (U/I). CSNK2A inhibitor **2** (10 μM) produced a 1.5-2.0 log reduction in virus compared vehicle control (Veh). Remdesivir (Rem, 2.5 μM) was included as a comparator. (C) Dose response effect of CSNK2A inhibitor **2** in primary HAE cells infected with SARS-CoV-2-nLuc (open circles) without affecting cell viability determined by LDH assay (closed circles), Remdesivir (✱, 2.5 μM) was included as a comparator. Assay performed in triplicate with data ranges shown by error bars.

### CSNK2A inhibitors block SARS-CoV-2 spike protein uptake

The broad substrate specificity of CSNK2 provides little guidance on the mechanism of antiviral activity of CSNK2A inhibitors.^30^ There are several key steps (entry, replication, packaging, and egress) where CSNK2A inhibition may impact in the virus lifecycle^1^ and over 40 different viral proteins have been shown to be CSNK2 substrates.^31^ Notably, several other host kinases have been recently implicated in the regulation of virus entry into cells.^32^ β-CoV infect cells following the attachment of their spike glycoprotein to receptors on the cell surface membrane.^33^ The primary mechanism by which the β-CoV-spike-membrane complex enters cells is a process of clathrin-mediated endocytosis (CME).^34^ The internalized β-CoV accumulates in the endolysosome until the action of host cell proteases leads to release of the virus mRNA. To explore whether CSNK2 inhibition could affect this critical step of β-CoV entry, we utilized an assay that measured the uptake of the SARS-CoV-2 spike protein trimer by CME into cells.^35^ His6-tagged spike protein was incubated with HEK293T-ACE2 cells for 30 min at 4°C to allow complex formation between the spike protein and ACE2, and at 37°C for 30 min to promote internalization by CME. The cells were then acid washed to remove extracellular spike protein and fixed. The intracellular spike protein was visualized using a His6 antibody and quantified by imaging (Figure 6). Vehicle treated cells showed efficient uptake of the His6-tagged SARS-CoV-2 spike protein. Treatment of the HEK293T-ACE2 cells with 1 μM of the CSNK2A chemical probe **SGC-CK2-1** or CSNK2A inhibitor **2** resulted in a 70–80% decrease in spike protein uptake (Figure 6). Notably, the negative control analog **SGC-CK2-1N** had no effect on spike protein uptake into the cells.

**Figure 6.**
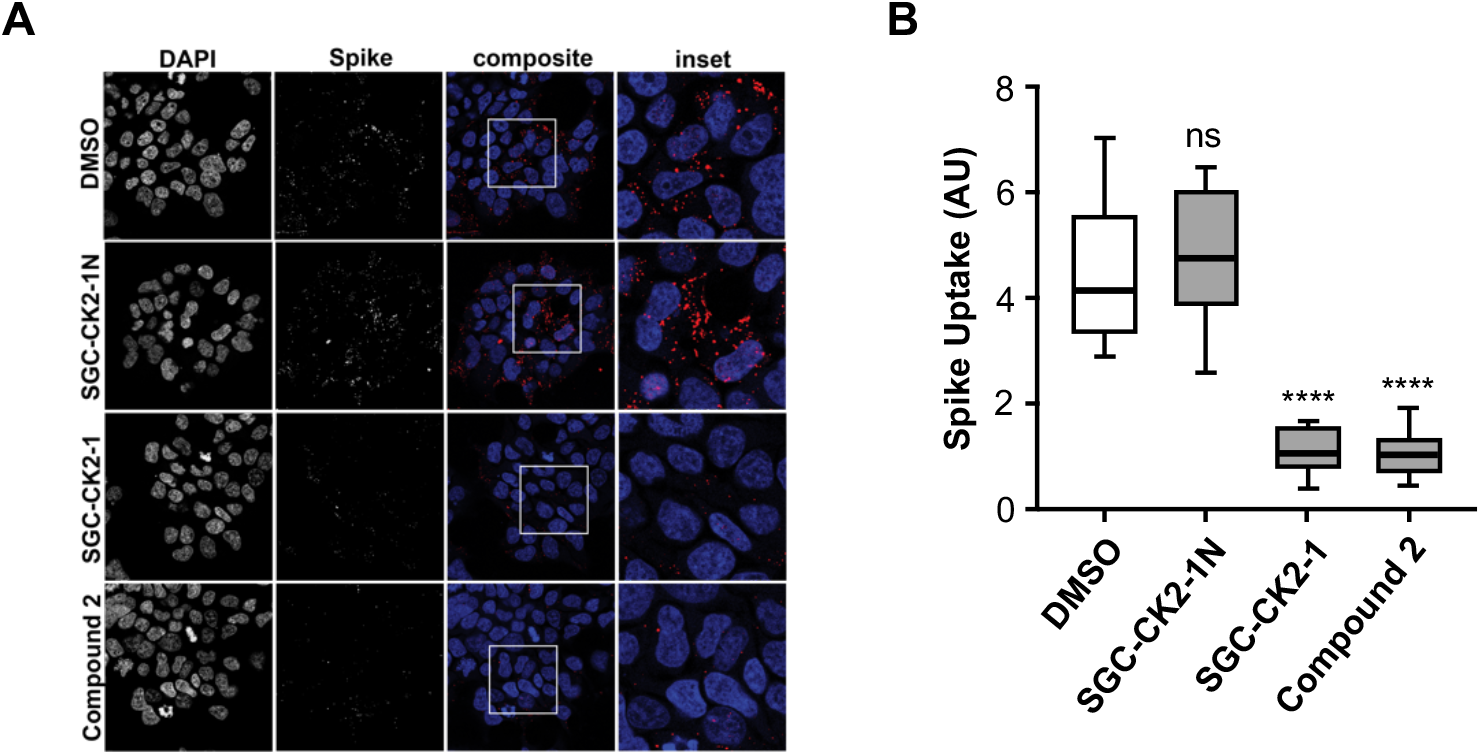
Effect of CSNK2A inhibition on His6-SARS-CoV-2 spike protein uptake into HEK293T-ACE2 cells. (A) Cell nuclei stained with DAPI. Spike protein detected using a His6 antibody, Cells were treated with 1 μM of CSNK2A inhibitors (**SGC-CK2-1** or Compound **2**), negative control (**SGC-CK2-1N**), or vehicle control (DMSO). (B) Quantification of the His6-SARS-CoV-2 spike protein uptake. The data is from 3 separate experiments with n = 9 for each sample. The total spike-His tag fluorescence was divided by the number of cells to control for the number of cells per frame. AU, arbitrary units; ns, not significant; **** p < 0.0001.

The proteomic and phosphoproteomic studies that identified a key role of CSNK2 in SARS-CoV-2 infection had focused on a role of the kinase in virus egress through remodeling of the extracellular matrix.^18^ Our data demonstrate that CSNK2 may also be involved in virus entry through CME (Figure 5), suggesting that β-CoV utilize a common kinase for multiple steps in viral trafficking during their life cycle. The Numb-associated kinases AAK1 and GAK have also been implicated in regulation of virus entry by CME.^32, 36^ However, inhibitors of these kinases generally demonstrate antiviral activity at only micromolar concentrations^12^ and the antiviral activity often does not track with kinase inhibition.^37, 38^ Furthermore, AAK1 inhibition was recently reported to block SARS-CoV-2 virus uptake only in a subset of cells that lack the ACE2 receptor,^39^ implicating a mechanism independent of CME. In our assays, selective chemical probes for AAK1^40^ or GAK^41^ failed to prevent β-CoV replication when used at their recommended 1 μM dose (SI Table 1). In contrast, we saw antiviral activity that tracked with CSNK2A potency (Figure 3) and robust inhibition of SARS-CoV-2 spike protein uptake by selective CSNK2A inhibitors (Figure 5).

### CSNK2 is a host target for inhibition of β-CoV

Multiple observations argue in favor of CSNK2A inhibition as an antiviral mechanism in the MHV replication assay. First, the high-quality chemical probe **SGC-CK2-1** showed inhibition of virus replication at nanomolar doses where it has remarkably high kinase selectivity.^24^ Second, the structurally related negative control analog **SGC-CK2-IN** had no effect on virus replication at does up to 100-fold higher in concentration. Third, silmitasertib, a chemotype of CSNK2A inhibitor with different chemical and physical properties also inhibited virus replication. Fourth, across three series of ATP-competitive 3-cyano-pyrazolo[1,5-*a*]pyrimidine CSNK2A inhibitors with substitutions at multiple sites on the heterocyclic core, the structure-activity for virus inhibition matched the potency of kinase inhibition (Figure 3). The combined chemogenomic evidence strongly implicates CSNK2A inhibition as the molecular mechanism of action of antiviral activity. Confirmation that CSNK2 is a host cell kinase required for β-CoV replication was provided by genetic knockdown of the essential regulatory subunit CSNK2B or dual knockdown of the catalytic subunits CSNK2A1 and CSNK2A2 (Figure 4D). While further studies will be required to dissect the molecular details of the signaling pathway that requires CSNK2 for virus uptake and its relative contribution to β-CoV replication, the potent anti-β-CoV activity of CSNK2A inhibition suggests that it may be a viable broad spectrum antiviral therapy for current and future zoonotic diseases at doses that could be attainable in clinical setting.

## ASSOCIATED CONTENT

### Supporting Information

**SI Figure 1 –** Cell viability determined by LDH assay in DBT cells after 10 h.

**SI Figure 2** – Comparison of CSNK2A1 and CSNK2A2 potency versus MHV inhibition

**SI Figure 3** – Human and mouse CSNK2A subunits show high sequence identity.

**SI Table 1** – SGC Kinase Chemical Probes tested in the MHV-nLuc assay in DBT cells

**Biology Methods** – Biological assays and reagents

**Chemistry Methods** – Synthesis and characterization of compounds (**8**–**33**)

## ACKNOWLEDGEMENT

Constructs for nanoBRET measurements of CSNK2A1 and CSNK2A2, were provided by Promega. CSNK2 antibodies were generously provided by Dr. David Litchfield (University of Western Ontario). We thank Aled Edwards and Kumar Saikatendu Singh for facilitating collaborative interactions, constructive criticism throughout the project, and for critical reading of the manuscript. Wuxi AppTec (Shanghai) and ChemSpace LLC provided chemical synthesis support. We acknowledge support of the Neuro Microscopy Imaging Centre and Advanced BioImaging Facility at McGill University. We thank Dr Jesse D. Bloom (University of Washington, Seattle) for the HEK-293T-ACE2 stable cell line.

