## Supporting Information for "Host kinase CSNK2 is a target for inhibition of pathogenic β-coronaviruses including SARS-CoV-2"

### Table of Contents

P1–4 **SI Figure 1** – Cell viability determined by LDH assay in DBT cells after 10 h.

P5 **SI Figure 2** – Comparison of CSNK2A1 and CSNK2A2 potency versus MHV inhibition

P6 **SI Figure 3** – Human and mouse CSNK2A subunits show high sequence identity.

P7 **SI Table 1** – SGC Kinase Chemical Probes tested in the MHV-nLuc assay in DBT cells

P8–12 **Biology Methods** – Biological assays and reagent**s**

P13–36 **Chemistry Methods** – Synthesis and characterization of compounds (**8–33**)

P39 **References**

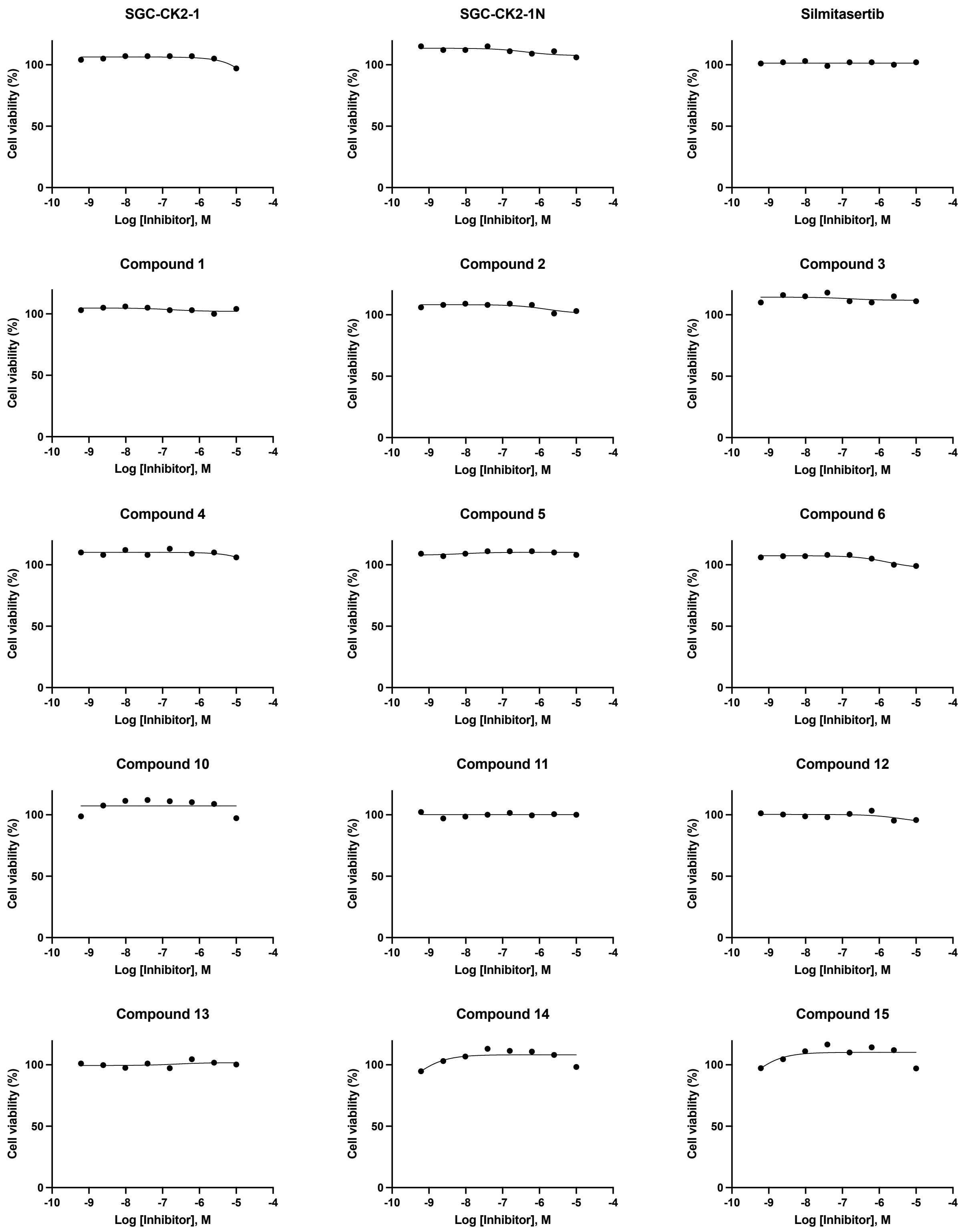
SGC-CK2-1
SGC-CK2-1N
Silmitasertib
Compound 1
Compound 2
Compound 3
Compound 4
Compound 5
Compound 6
Compound 10
Compound 11
Compound 12
Compound 13
Compound 14
Compound 15
Cell viability (%)
Log [Inhibitor], M

Compound 16

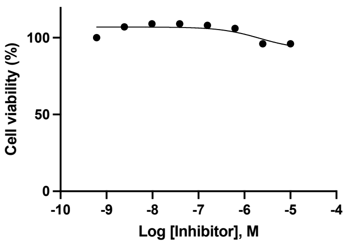


Compound 17

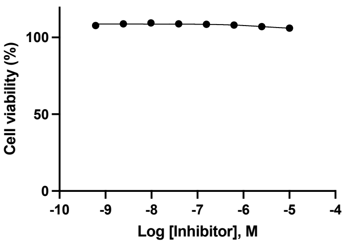


Compound 18

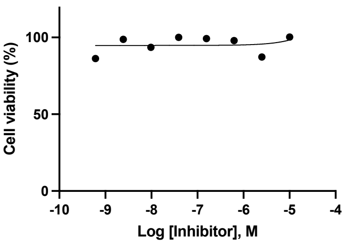


Compound 19

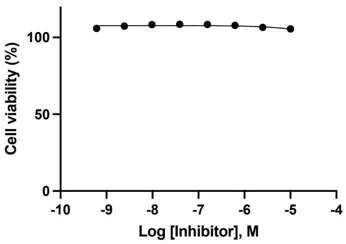


Compound 20

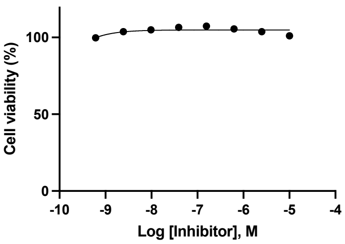


Compound 21

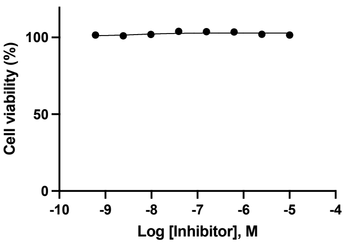


Compound 22

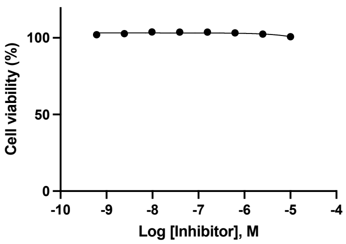


Compound 23

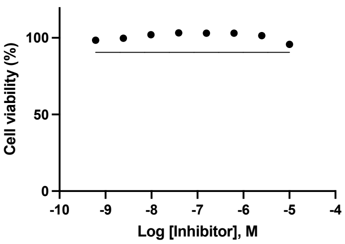


Compound 24

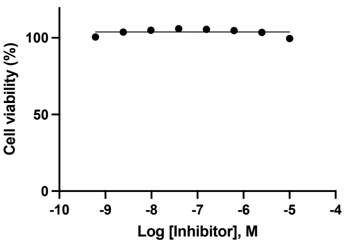


Compound 25

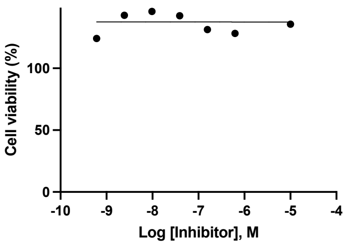


Compound 26

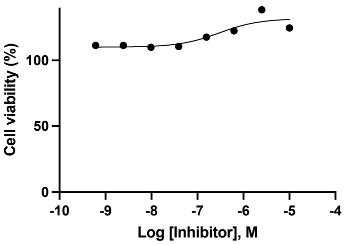


Compound 27

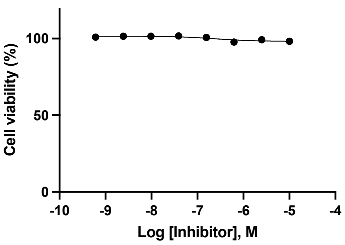


Compound 28

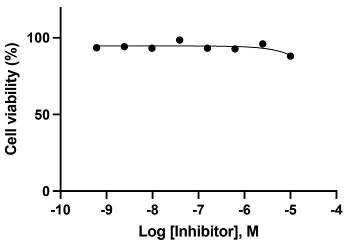


Compound 29

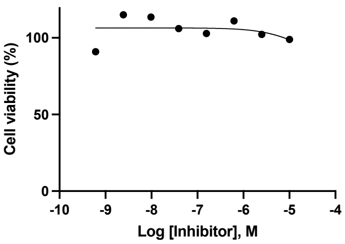


Compound 30

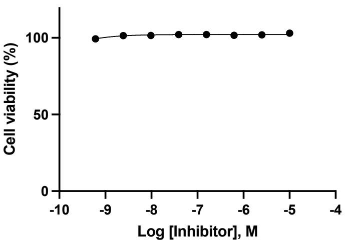

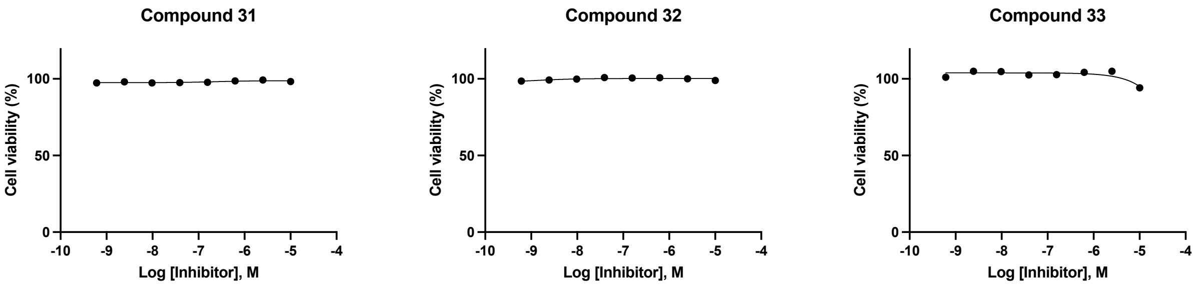


**SI Figure 1.** Cell viability determined by LDH assay in DBT cells after 10 h.

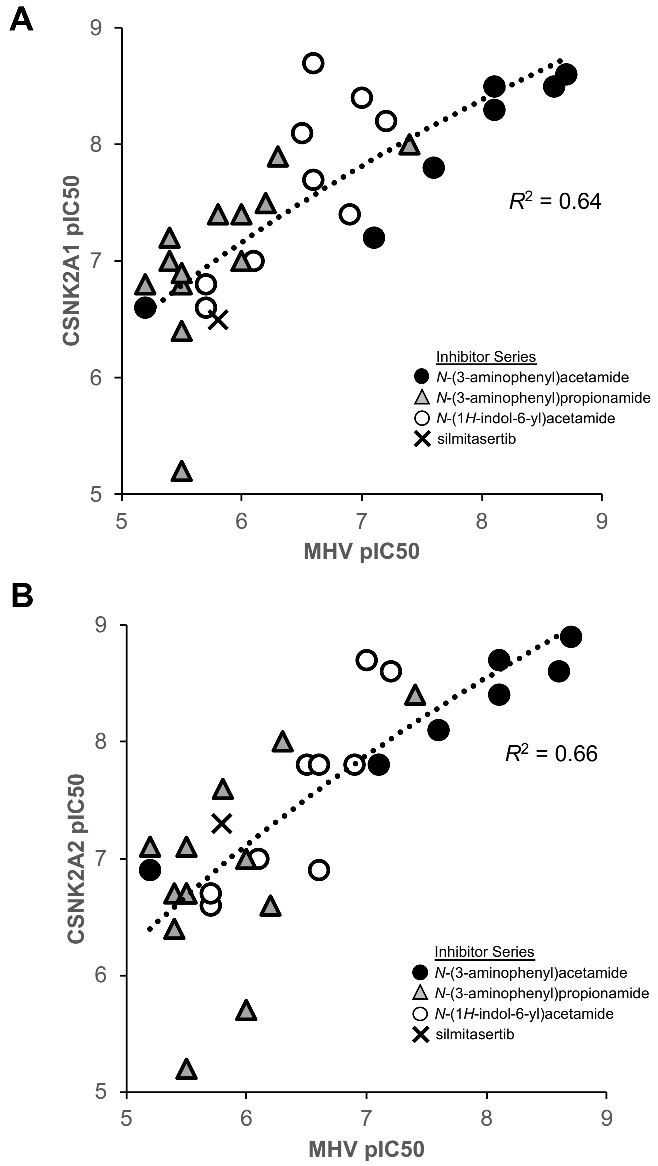


**SI Figure 2.** Correlation of potency for CSNK2A target engagement with inhibition of β-CoV replication across three subseries of pyrazolo[1,5-*a*]pyrimidines and silmitasertib. (A) CSNK2A1. with an $R^2$ = 0.64. (B) CSNK2A2 with an $R^2$ = 0.66. Inactive analogs **SGC-CK2-1N**, **20**, **21**, **22**, **31**, and **32** were not included in the analysis.

```
CLUSTAL O(1.2.4) multiple sequence alignment


sp|P68400|CSNK2A1_HUMAN      MSGP-VPSRARVYTDVNTHRPREYWDYESHVVEWGNQDDYQLVRKLGRGKYSEVFEAINI   59
sp|Q60737|CSNK2A1_MOUSE      MSGP-VPSRARVYTDVNTHRPREYWDYESHVVEWGNQDDYQLVRKLGRGKYSEVFEAINI   59
sp|P19784|CSNK2A2_HUMAN      MPGPAAGSRARVYAEVNSLRSREYWDYEAHVPSWGNQDDYQLVRKLGRGKYSEVFEAINI   60
sp|O54833|CSNK2A2_MOUSE      MPGPAAGSRARVYAEVNSLRSREYWDYEAHVPSWGNQDDYQLVRKLGRGKYSEVFEAINI   60
                             * ** . ******::**: * *******:** .***************************

sp|P68400|CSNK2A1_HUMAN      TNNEKVVVKILKPVKKKKIKREIKILENLRGGPNIITLADIVKDPVSRTPALVFEHVNNT   119
sp|Q60737|CSNK2A1_MOUSE      TNNEKVVVKILKPVKKKKIKREIKILENLRGGPNIITLADIVKDPVSRTPALVFEHVNNT   119
sp|P19784|CSNK2A2_HUMAN      TNNERVVVKILKPVKKKKIKREVKILENLRGGTNIIKLIDTVKDPVSKTPALVFEYINNT   120
sp|O54833|CSNK2A2_MOUSE      TNNERVVVKILKPVKKKKIKREVKILENLRGGTNIIKLIDTVKDPVSKTPALVFEYINNT   120
                             ****:*****************:********* ***.* * ******:*******::***

sp|P68400|CSNK2A1_HUMAN      DFKQLYQTLTDYDIRFYMYEILKALDYCHSMGIMHRDVKPHNVMIDHEHRKLRLIDWGLA   179
sp|Q60737|CSNK2A1_MOUSE      DFKQLYQTLTDYDIRFYMYEILKALDYCHSMGIMHRDVKPHNVMIDHEHRKLRLIDWGLA   179
sp|P19784|CSNK2A2_HUMAN      DFKQLYQILTDFDIRFYMYELLKALDYCHSKGIMHRDVKPHNVMIDHQQKKLRLIDWGLA   180
sp|O54833|CSNK2A2_MOUSE      DFKQLYQILTDFDIRFYMYELLKALDYCHSKGIMHRDVKPHNVMIDHQQKKLRLIDWGLA   180
                             ******* ***:********:********* ****************:::**********

sp|P68400|CSNK2A1_HUMAN      EFYHPGQEYNVRVASRYFKGPELLVDYQMYDYSLDMWSLGCMLASMIFRKEPFFHGHDNY   239
sp|Q60737|CSNK2A1_MOUSE      EFYHPGQEYNVRVASRYFKGPELLVDYQMYDYSLDMWSLGCMLASMIFRKEPFFHGHDNY   239
sp|P19784|CSNK2A2_HUMAN      EFYHPAQEYNVRVASRYFKGPELLVDYQMYDYSLDMWSLGCMLASMIFRREPFFHGQDNY   240
sp|O54833|CSNK2A2_MOUSE      EFYHPAQEYNVRVASRYFKGPELLVDYQMYDYSLDMWSLGCMLASMIFRKEPFFHGQDNY   240
                             *****.*******************************************:******:***

sp|P68400|CSNK2A1_HUMAN      DQLVRIAKVLGTEDLYDYIDKYNIELDPRFNDILGRHSRKRWERFVHSENQHLVSPEALD   299
sp|Q60737|CSNK2A1_MOUSE      DQLVRIAKVLGTEDLYDYIDKYNIELDPRFNDILGRHSRKRWERFVHSENQHLVSPEALD   299
sp|P19784|CSNK2A2_HUMAN      DQLVRIAKVLGTEELYGYLKKYHIDLDPHFNDILGQHSRKRWENFIHSENRHLVSPEALD   300
sp|O54833|CSNK2A2_MOUSE      DQLVRIAKVLGTDELYGYLKKYHIDLDPHFNDILGQHSRKRWENFIHSENRHLVSPEALD   300
                             ************::**.*:.**:*:***:******:*******.*:****:*********

sp|P68400|CSNK2A1_HUMAN      FLDKLLRYDHQSRLTAREAMEHPYFYTVVKDQARMGSSSMPGGSTPVSSANMMSGISSVP   359
sp|Q60737|CSNK2A1_MOUSE      FLDKLLRYDHQSRLTAREAMEHPYFYTVVKDQARMSSTSMAGGSTPVSSANMMSGISSVP   359
sp|P19784|CSNK2A2_HUMAN      LLDKLLRYDHQQRLTAKEAMEHPYFYPVVKEQSQPCADNAV----------LSSGLTAAR   350
sp|O54833|CSNK2A2_MOUSE      LLDKLLRYDHQQRLTAKEAMEHPYFYPVVKEQSQPCAENTV----------LSSGLTAAR   350
                             :**********.****:********* ***:*::  : .            : **:::.


sp|P68400|CSNK2A1_HUMAN      TPSPLGPLAGSPVIAAANPLGMPVPAAAGAQQ   391
sp|Q60737|CSNK2A1_MOUSE      TPSPLGPLAGSPVIAAANSLGIPVPAAAGAQQ   391
sp|P19784|CSNK2A2_HUMAN      --------------------------------   350
sp|O54833|CSNK2A2_MOUSE      --------------------------------   350
```

**SI Figure 3.** Human and mouse CSNK2A subunits show high sequence identity. Multiple sequence alignment was performed by Clustal Q. The crystallographically-defined kinase domain is marked by the grey bar. Human and mouse CSNK2A1 are identical within the kinase domain. Only a single amino acid (shaded in black) in is different between the human and mouse CSNK2A2 within the kinase domain. CSNK2A2 E253D is at the base of the C-lobe >30Å from the ATP-binding pocket. Amino acid residues with sidechains within 4Å of pyrazolo[1,5-a]pyrimidine **4** in the cocrystal structure with human CSNK2A1 (PDB 5H8G) are marked with black dots. 17/17 contact residues are identical between the human and mouse sequences. 15/17 inhibitor contact residues are identical across CSNK2A1 and CSNK2A2. The 2 residues that differ (CSNK2A1 H115Y and V116I) are underlined.

| Kinase | Chemical Probe (1 µM) | MHV (%I) | Negative Control Analog (1 µM) | MHV (%I) |
|---|---|---|---|---|
| **AAK1** | **SGC-AAK1-1** | 28 | | |
| ACVR1B | TP-008 | 28 | AI11 | 5 |
| AKT1/2/3 | MK-2206 | 70 | | |
| AURA | MK-5108 | 9 | | |
| AURA | Alisertib | 16 | | |
| AURA/B | PF-03814735 | 58 | | |
| AURB | Barasertib | -8 | | |
| AXL | TP-0903 | 30 | | |
| AXL/MER | LDC1267 | 19 | | |
| AXL/MET | NPS-1034 | 15 | | |
| BRAF | GDC-0879 | 18 | | |
| BTK | CGI 1746 | 19 | | |
| CAMK2 | KN-93 | 20 | | |
| CAMK2 | KN-62 | 28 | | |
| CAMKK1/2 | STO-609 | 6 | | |
| CAMKK1/2 | SGC-CAMKK2-1 | 15 | SGC-CAMKK2-1N | 66 |
| CASK | NR162 | 40 | NR187 | 26 |
| CDK8/19 | BI-1347 | 41 | BI-1374 | 44 |
| CDK9 | LDC000067 | -17 | | |
| CLK1/2/3/4 | T3-CLK | -4 | T3-CLK-N | -1 |
| CLK1/2/4 | SGC-CLK-1 | 61 | SGC-CLK-1N | 1 |
| CLK1/2/4 | MU1210 | 2 | MU140 | 9 |
| CLK1/4 | KH-CB19 | -15 | | |
| DDR1/2 | DDR-TRK-1 | 16 | DDR-TRK-1N | 5 |
| DDR1/2 | SR302 | -11 | SR301 | 0 |
| EGFR | Afatinib | 51 | | |
| EPHB4 | NVP-BHG712 | 57 | | |
| ERK5 | XMD8-92 | 9 | | |
| **GAK** | **SGC-GAK-1** | 4 | SGC-GAK-1N | 38 |
| GSK3A/B | CHIR-99021 | 12 | | |
| IGF1R | Picropodophyllin | 44 | | |
| JAK3 | FM-381 | 36 | FM-479 | 30 |
| LIMK1/2 | TH-257 | 10 | TH-263 | 31 |
| LRRK2 | MLi-2 | 37 | Mli-2-NC | 20 |
| MAPK1/3 | ERKi | 27 | ERKi-NC | 25 |
| MAPK7 | BAY-885 | 23 | BAY-693 | 16 |
| MAPK14 | SR318 | 13 | | |
| MAPK14 | VX-702 | 15 | | |
| MAPK14 | L-Skepinone | 19 | FM-743 | -11 |
| MAPK14 | FS-694 | 0 | | |
| MAPKAP | PF-3644022 | 34 | | |
| MEK1/2 | Trametinib | 1 | | |
| MEK1/2 | Selumetinib | 11 | | |
| MER | UNC2250 | 32 | | |
| MET | SGX-523 | -2 | | |
| MET | BAY-474 | 4 | BAY-827 | 0 |
| MET | Tepotinib | 27 | | |
| PAK1 | NVS-PAK-1 | -32 | NVS-PAK1-C | 10 |
| PAK1/2/3 | FRAX486 | 47 | | |
| PDK1 | GSK2334470 | 35 | | |
| PKCA/B/G | BisindolyImaleimid | 37 | | |
| PKCB | Enzastaurin | 37 | | |
| PRKAA1 | BAY-3827 | -4 | BAY-974 | -7 |
| PTK2 | PF-562271 | 34 | | |
| PTK2 | PF-04554878 | 19 | PF-00911705 | 23 |
| PTK2 | Defacitinib | 38 | | |
| PTK2 | PND-1186 | 26 | | |
| RAF1 | GW5074 | 35 | | |
| RIPK1 | TP_030_1 | -1 | TP_030_2 | 44 |
| ROCK1/2 | BAY-549 | 51 | BAY-4900 | -8 |
| ROCK1/2 | RKI-1447 | 13 | | |
| ROCK1/2 | Y-27632 | 11 | | |
| ROCK1/2 | GSK429286A | 18 | | |
| S6K1 | PF-4708671 | 0 | | |
| SGK1/2 | GSK650394 | 28 | | |
| STK17B | SGC-STK17B-1 | -36 | SGC-STK17B-1N | -48 |
| SYK | MRL-SYKi | -23 | MRL-SYKi-NC | 25 |
| SYK | PRT062607 | 40 | | |
| TBK1/IKBKE | BAY-985 | 23 | BAY-440 | 8 |
| TIE1/2 | BAY-826 | 42 | BAY-309 | 46 |
| TRKA/B/C | GNF-5837 | 51 | | |

**SI Table 1.** SGC Kinase Chemical Probes tested in the MHV-nLuc assay in DBT cells. All chemical probes and their corresponding negative control analogs (when available) were tested at 1 µM for inhibition of MHV-nLuc replication as described in the Methods. Results are displayed as the percentage of inhibition of luciferase compared to vehicle (DMSO) colored coded by relative inhibition from green (no inhibition) to red (70% inhibition). For each inhibitor the target kinase is indicated. For several kinases multiple chemical probes representing different chemotypes were tested. Chemical probes with >50% inhibition of MHV-nLuc replication were scored as active. Chemical probes for AAK1 (SGC-AAK-1) and GAK (SGC-GAK-1) highlighted in bold on a gray background were inactive.

### BIOLOGY METHODS

**Cell Culture.** DBT cells were cultured at 37 °C in Dulbecco's modified Eagle medium (DMEM; Sigma) supplemented with 10% fetal bovine serum (Gibco) and penicillin and streptomycin (Sigma).

Primary human airway epithelial (HAE) cells were cultured according to standard protocol.[1] Briefly, HAE cells were expanded on plates coated with Bovine Collagen Type I/III (Advanced BioMatrix) and cultured in BEGM media. HAE cells were seeded onto transwells coated with HPC Collagen IV (Sigma) and cultured with ALI media. Cells were visually inspected for hallmarks of differentiation and used in studies between days 28-35 post seeding into transwells.

HEK-293 cells were cultured in DMEM supplemented with 10% FBS. Cells were incubated in 5% $CO_2$ at 37 °C. Cells lines were passaged every 72 h with trypsin and not allowed to reach confluency.

**NanoBRET Assay.** Assays were run as previously described.[2] Briefly, a 10 µg/mL solution of DNA in Opti-MEM without serum was made containing 9 µg/mL of Carrier DNA (Promega) and 1 µg/mL of CSNK2A1-NL or CSNK2A2-NL (Promega) for a total volume of 1.05 mL. 31.5 µL of FuGENE HD (Promega) was added to form a lipid:DNA complex. The solution was then mixed by inversion 8 times and incubated at room temperature for 20 min. The transfection complex (1.082 mL) was then gently mixed with 21 mL of HEK-293 cells (ATCC) suspended at a density of 2 x $10^5$ cells/mL in DMEM (Gibco) + 10% FBS (Corning). 100 µL was dispensed into 96-well tissue culture treated plates (Corning #3917) and incubated at 37 °C in 5 % $CO_2$ for 24 h. The media was removed and replaced with 85 µL of Opti-MEM without phenol red. A total of 5 µL per well of 20 µM nanoBRET Tracer K5 in Tracer Dilution Buffer (Promega N291B)) was added to all wells, except the "no tracer" control wells. Test compounds (10 mM in DMSO) were diluted in Opti-MEM media (99%) to prepare 1% DMSO stock solutions and evaluated at eleven concentrations. A total of 10 µL per well of the 10-fold test compound stock solutions (final assay concentration of 0.1% DMSO) were added. For "no compound" and "no tracer" control wells, a total of 10 µL per well of OptiMEM plus DMSO (9 µL Opti-MEM with 1 µL DMSO) was added for a final concentration of 1% DMSO. 96-well plates containing cells with nanoBRET Tracer K5 and test compounds (100 µL total volume per well) were equilibrated (37 °C / 5 % $CO_2$) for 2 h. The plates were cooled to room temperature for 15 min. nanoBRET NanoGlo substrate (Promega) at a ratio of 1:166 to Opti-MEM media in combination with extracellular NanoLuc Inhibitor (Promega) diluted 1:500 (10 µL of 30 mM stock per 5 mL Opti-MEM plus substrate) were combined to create a 3X stock solution. A total of 50 µL of the 3X substrate/extracellular NL inhibitor were added to each well. The plates were read within 10 min on a GloMax Discover luminometer (Promega) equipped with 450 nm BP filter (donor) and 600 nm LP filter (acceptor) using 0.3 s integration time. Raw milliBRET (mBRET) values were obtained by dividing the acceptor emission values (600 nm) by the donor emission values (450 nm) and multiplying by 1000. Averaged control values were used to represent complete inhibition (no tracer control: Opti-MEM + DMSO only) and no inhibition (tracer only control: no compound, Opti-MEM + DMSO + Tracer K5 only) and were plotted alongside the raw mBRET values. The data with n=3 biological replicates was first normalized and then fit using Sigmoidal, 4PL binding curve in Prism Software to determine $IC_{50}$ values.

**Viruses.** *MHV-nLuc:* The MHV-A59 G plasmid was engineered to replace most of the coding sequence for orf4a and 4b with nLuc. Briefly, nucleotides 27,983 to 28,267 were removed and replaced with SalI and SacII restriction sites; approximately 111 bp of the 3' end of orf4B was left to maintain the TRS for orf5. nLuc was PCR amplified with primers 5'nLuc SalI (5'-NNNNNNGTCGACATGGTCTTCACACTCGAAGATTTC-3') and 3'nLuc SacII (5'-NNNNNNCCGCGGTTACGCCAGAATGCGTTCGCAC-3'), digested with SalI and SacII and then cloned into the G plasmid which had been similarly digested. A sequence verified G-nLuc plasmid was used with MHV-A59 wild type A, B, C, D, E and F plasmids to recover virus expressing nLuc and the recombinant virus sequence verified. MHV-nLuc virus stocks were grown on DBT cells and their titers were determined using the 50% tissue culture infectious dose ($TCID_{50}$) assay.

*SARS-CoV2-nLuc:* A549-ACE2 cells (85-95% confluent) were infected at MOI of 0.01 with ic-SARS-CoV-2-nluc[3] in DMEM containing 5% heat-inactivated serum. Infected monolayers were incubated at 37°C with 5% $CO_2$ until CPE involved approximately 50% of the monolayer (generally between 66 and 72 h). Infected cell culture supernatant was recovered and clarified by centrifugation and aliquots of the clarified supernatant were frozen at -80 °C until use.

**MHV Assay.** DBT cells were plated in 96 well plates to be 80% confluent at the start of the assay. Test compounds were diluted to 15 µM in DMEM. Serial 4-fold dilutions were made in DMEM, providing a concentration range of 15 µM to 0.22 µM. Media was aspirated from the DBT cells and 100 µL of the diluted test compounds were added to the cells for 1 h at 37 °C. After 1 h, MHV-nLuc was added at an MOI of 0.1 in 50 µL DMEM so that the final concentration of the first dilution of compound was 10 µM (T=0). After 10 h, the media was aspirated, and the cells were washed with PBS and lysed with passive lysis buffer (Promega) for 20 min at room temperature. Relative light units (RLUs) were measured using a luminometer (Promega; GloMax). Triplicate data was analyzed in Prism Graphpad to generate $IC_{50}$ values.

**HAE Assay.** HAE cultures were washed 3 times with pre-warmed PBS (20 min each wash) to remove mucus from the apical surface. After the last apical wash, spent ALI media was removed and replaced with media containing drug, DMSO or media only as needed. Immediately after the media was replaced, 100 μL of ic-SARS-CoV-2-nluc[3] was added to the apical side of the HAE cultures to achieve MOI = 0.5. Cultures were returned to the incubator and allowed to infect for 2 h. The inoculum was removed and the apical surface was washed 3 times with PBS to remove unbound virus before the cells were returned to the incubator. 24 h post-infection, the cells were washed by adding 100 µL of pre-warmed PBS to the apical surface and incubating at 37°C for 20 min. The apical wash was removed, the inserts transferred to a new 12 well plate and 150 µL of Passive Lysis Buffer (Promega) added to each well. After 10 min incubation at room temperature, the inserts were scraped with a pipet tip and the lysed cell mixture was recovered. Fifty microliter aliquots of lysed cell mixture was transferred to a clear bottom, black-walled plate and mixed with 50 µL of NanoGlo Reagent (Promega). Luminescence was read on a GloMax instrument (Promega). For calculations, wells containing passive lysis buffer mixed with NanoGlo Reagent were used as background luminescence and this background was subtracted from the RLU of each sample. RLU (background adjusted) were graphed directly in the bar graphs. For dose response curves, the percent inhibition was calculated as follows: (1-(Sample RLU-background)/(virus only RLU-background adjusted)) x 100 with range normalized from 0-100 and $IC_{50}$ calculated using GraphPad Prism.

**LDH Assay.** DBT cells were plated to be 80 % confluent at the start of the assay. Compounds were diluted as for the MHV assay and incubated with cells at 37 °C for 1 h. After 1 h, 50 µL of DMEM was added to the cells (T=0). 45 minutes before harvest, lysis buffer was added to positive wells. LDH activity in cell-free supernatants was measured at 10 h after infection using the Sigma Tox7 kit as per the manufacturer’s directions.

**siRNA Knockdown.** SMARTPool ON-TARGETplus mouse siRNAs were purchased for Csnk2a1 (L-058653-00-0005), Csnk2a2 (L-051582-00-0005), Csnk2b (L-049417-00-0005), or non-targeting (D-001810-10-05) genes (Horizon). 200 µL of transfection master mix (12.5 nmol siRNA, RNAi Max, OptiMEM) was reverse transfected with DBT cells and incubated at 37 °C for 48 h. Cells were either collected for western analysis or trypsionized and replated with fresh siRNA transfection master mix. Replated cells were 80 % confluent and used for MHV assay experiments.

**qRT-PCR.** Cells were scraped, pelleted, and stored at -80 °C until time of analysis. RNA was extracted from cell pellets using TRIzol (ThermoFisher Scientific) and chloroform. After a 10 min spin, an equal volume of isopropanol was added to the aqueous layer and RNA was precipitated overnight at -20 °C. RNA was washed with ethanol and DNAse treated (TURBO DNase, ThermoFisher Scientific). RNA was quantified by NanoDrop (ThermoFisher Scientific) and 2 µg of RNA was used to make cDNA (High Capacity cDNA Reverse Transcription kit, Thermo Fisher Scientific). For real-time PCR, 0.5 µM gene-specific primers (csnk2a1: Fwd GGTGAGGATAGCCAAGGTTCTG, Rev TCACTGTGGACAAAGCGTTCCC; csn2a2: Fwd GGATTACTGCCACAGCAAGGGA, Rev GGATGATAGAACTCTGCCAGACC; csnk2b: Fwd CAGAGCGACTTGATCGAACAGG, Rev CGAGGACAGTAGCCAAAGTCTC) and 1X SYBR green master mix was added to 2 µL of cDNA. RNA abundance was quantified using a standard curve generated from 10-fold serial dilutions of a DNA standard specific for each primer pair. The relative expression at t = 4, 8, and 12 h post-infection was determined by dividing the RNA abundance at each timepoint by the value determined following mock infection (t = 0).

**Western Blot Analysis.** Cells were scraped and pelleted for western blot analysis and stored at -80 °C until time of analysis. Pellets were thawed on ice and lysed for 10 minutes in radioimmunoprecipitation assay buffer (RIPA: 50 mM Tris-HCl [pH 7.4], 150 mM NaCl, 1 mM EDTA, 1% NP-40, 1% sodium deoxycholate) supplemented with 1x Complete protease inhibitor cocktail (Roche). Cells were spun at 4 °C to pellet debris and protein concentration determined via Bradford assay (VWR). Equal amounts of protein were resolved on a 10 % SDS-PAGE gel and transferred to nitrocellulose membranes (Amersham). Membranes were blocked for 1 h at room temperature with 5% nonfat milk in TBS-T (20 mM Tris-HCl [pH 7.6], 140 mM NaCl, 0.1% Tween 20). Membranes were washed with TBS-T prior to incubation with primary antibody. Rabbit polyclonal antibodies were dissolved in 5% bovine serium albumin (BSA) in TBS-T and incubated overnight at 4 °C. Blots were washed twice in TBS-T for 10 min prior to incubation with secondary horseradish peroxidase-conjugated rabbit antibody for 1 h at room temperature. Blots were imaged using chemiluminescent digital imager (Bio-Rad). Antibodies were provided by Dr. David Litchfield (Western University) and have been described previously:[4-6] anti-CSNK2A1 (KLH-CK2$\alpha$; 1:5000), anti-CSNK2B antibody (KLH-CK2$\beta$; 1:10000), anti-CSNK2A1/CSNK2A2 antibody (1:2000).

**Spike Uptake Assay.** The protocol used for the uptake of spike protein has been described previously.[7] Briefly, HEK293T-ACE2 cells were seeded onto poly-L-lysine treated coverslips, 24 h prior to experimentation. 1 h prior to the addition of spike protein, cell media were changed to starvation media (lack of serum) along with 1 μM test compounds and DMSO (vehicle control). Spike protein (5 μg per well) was added to each coverslip, and cells were incubated on ice for 30 min. Cells were then washed with PBS, and the media was replaced with fresh starvation media supplemented with the same test compound at 1 μM. Cells were then incubated for 30 min at 37 °C. Prior to fixation, cells were acid washed for 60 sec, followed by an acid rinse, to remove any extracellular spike protein. This was followed by PBS wash and fixation for 10 min with PFA at 4 °C. Cells were then permeabilized and blocked with 5% Bovine Serum Albumin. His-tag antibody (HIS.H8) conjugated with Dylight 550 (Thermofisher) was used to identify spike protein uptake. Cells were then mounted and imaged using Leica SP8 microscope. Quantification was done with Leica LAS X software, with statistical calculations and graphs produced using Prism Graphpad software.

### CHEMISTRY METHODS

#### Synthesis of Intermediates (37–44)

*N-(2-fluoro-5-nitrophenyl)propionamide (**37**):* To a solution of 2-fluoro-5-nitroaniline (**36**) (5.00 g,

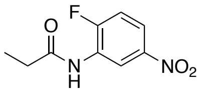

32.03 mmol) in DCM (60 mL) was added TEA (9.72 g, 96.08 mmol). Then the reaction mixture was cooled down to 0℃, and propionyl chloride (3.26 g, 35.23 mmol) was added to the mixture dropwise and was stirred at 25°C. After 3 hours, the reaction mixture was quenched by water (20 mL) and extracted with EtOAc (100 mL×2). The combined organic layers were dried over $Na_2SO_4$, filtered, and concentrated in *vacuo*. The crude product was then purified by flash chromatography (silica gel) eluting with a gradient of petroleum ether: EtOAc (100:0 to 80:20) to afford 2.90 g (32.2%) of the title compound **37** as a light-yellow liquid. $^1$H NMR (400 MHz, DMSO-$d_6$) $\delta$ H = 10.10 (s, 1H), 9.00 (dd, *J* = 2.8, 6.8 Hz, 1H), 8.047.98 (m, 1H), 7.55 (t, *J* = 10.0 Hz, 3H), 2.48−2.41 (m, 2H), 1.08 (t, J = 7.6 Hz, 3H). LCMS $R_t$ = 0.473 min in 1 min chromatography, purity 75.5%, MS ESI *m/z*: 231.0 [M+H]$^+$.

*tert-Butyl (2-(benzylamino)ethyl)(ethyl)carbamate (**37b**):* To a solution of compound **37a** (1.00 g,

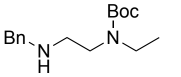

5.31 mmol) in MeOH (15 mL) was added benzaldehyde (564 mg, 5.31 mmol) slowly at 25°C. After addition, the mixture was stirred at 25°C for 1hour, and then $NaBH_4$ (603 mg, 15.93 mmol) was added at 0°C. The resulting mixture was stirred at 25°C for 2 hours under $N_2$ atmosphere. The reaction mixture was quenched in water (15 mL) and extracted with DCM (3 × 30 mL). The combined organic layer was washed with brine (2 × 6 mL), dried over $Na_2SO_4$, filtered, and concentrated to provide the crude product. The residue was purified by flash silica gel chromatography (eluent of 0%~3%, MeOH/DCM) to get **37b** (1.45 g, 61.5%) as a white oil. $^1$H NMR (400 MHz, DMSO-$d_6$) $\delta$ 7.51−7.14 (m, 5H), 3.69 (s, 2H), 3.25−3.10 (m, 4H), 2.58 (t, J = 6.8 Hz, 2H), 1.41−1.30 (m, 9H), 1.00 (t, *J* = 6.4 Hz, 3H). LCMS $R_t$ = 1.242 min in 2.5 min chromatography, purity 62.9%, MS ESI *m/z*: 279.2 [M+H]$^+$.

*tert-Butyl (2-(benzyl(methyl)amino)ethyl)(ethyl)carbamate (**37c**):* To a solution of compound **37b**

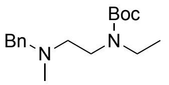

(1.40 g, 5.03 mmol) in DCM (15 mL) was added TEA (1.53 g, 15.09 mmol) and paraformaldehyde (453 mg, 15.09 mmol) at 25℃. The mixture was stirred at 25 °C for 0.5 hour. Then $NaBH_3CN$ (1.90 g, 30.17 mmol) was added to the mixture. The suspension was degassed and purged with $N_2$ three times. The mixture was stirred under $N_2$ at 25°C for 2 hours. The reaction mixture was quenched in water (10 mL) and extracted with EtOAc (3 × 20 mL). The combined organic layer was washed with brine (2 × 5 mL), dried over $Na_2SO_4$, filtered, and concentrated to provide the crude product. The residue was purified by flash silica gel chromatography (eluent of 0%~3%, MeOH/DCM) to get **37c** (1.85 g, 99.8%) as an off-white oil. $^{1}H$ NMR (400 MHz, DMSO-$d_6$) $\delta$ 7.33−7.20 (m, 5H), 3.49 (s, 2H), 3.30−3.20 (m, 2H), 3.17−3.10 (m, 2H), 2.46−2.40 (m, 2H), 2.23−2.09 (m, 3H), 1.40−1.30 (m, 9H), 1.05-0.95 (m, 3H). LCMS $R_t$ = 1.057 min in 2.5 min chromatography, purity 79.3%, MS ESI *m/z*: 293.2 $[M+H]^+$.

*tert-Butyl ethyl(2-(methylamino)ethyl)carbamate (**37d**):* To a solution of compound **37c** (1.80 g, 6.16mmol) in MeOH (20 mL) was added Pd/C (1.00 g, 10% purity) under $N_2$ atmosphere. The suspension was degassed and purged with $H_2$ three times. The reaction mixture was stirred under $H_2$ (15 Psi) at 25°C for 2 hours. The reaction mixture was filtered through a pad of Celite®, washed with MeOH (20 mL), and the combined organic extracts evaporated to dryness in *vacuo* to afford the crude product **37d** (579 mg, 97.2%) as a yellow solid, which was used to next step directly.

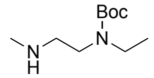


*tert-Butyl methyl(2-(methyl(4-nitro-2-propionamidophenyl)amino)ethyl)carbamate (**38a**):* To a solution of compound **37** (500 mg, 2.36 mmol) and tert-butyl methyl(2-(methylamino)ethyl)carbamate (532 mg, 2.83 mmol) in MeCN (8 mL) was added $K_2CO_3$ (977 mg, 7.07 mmol) at 25°C. Then the mixture was stirred at 100°C for 10 hours. The cooled black suspension was diluted with $H_2O$ (50 mL), extracted with EtOAc (3 × 50 mL), the combined organic extracts washed with brine (20 mL), dried over $Na_2SO_4$, and concentrated in *vacuo*. The crude product was then purified by flash chromatography (silica gel) eluting with a gradient of petroleum ether: EtOAc (100:0 to 70:30) to afford 718 mg (77.3%) of the title compound as a pale yellow liquid. $^{1}H$ NMR (400 MHz, DMSO-$d_6$) $\delta$ 9.30 (s, 1H), 8.49 (s, 1H), 7.92 (dd, *J* = 2.8, 8.8 Hz, 1H), 7.167.15 (m, 1H), 3.36−3.32 (m, 4H), 2.93 (s, 3H), 2.692.63 (m, 3H), 2.45 (br d, *J* = 7.2 Hz, 2H), 1.35 (s, 9H), 1.12 (t, *J* = 7.6 Hz, 3H). LCMS $R_t$ = 0.561 min in 4 min chromatography, purity 96.6%, MS ESI *m/z*: 381.1 $[M+H]^+$.

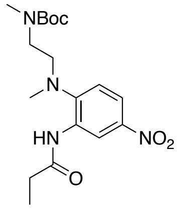


*tert-Butyl (2-(methyl(4-nitro-2-propionamidophenyl)amino)ethyl)carbamate (**38b**)*: Intermediate **38b** was prepared from **37** and *tert*-butyl (2-(methylamino)ethyl)carbamate in the same manner as described for the preparation of **38a**. Black brown oil (2.09 g, crude). $^{1}H$ NMR (400 MHz, DMSO-$d_6$) $\delta$ 9.20 (s, 1H), 8.55 (d, *J* = 2.4 Hz, 1H), 7.91 (dd, *J* = 2.8, 9.2 Hz, 1H), 7.19 (d, *J* = 9.2 Hz, 1H), 6.89 (s, 1H), 3.13 (br s, 4H), 2.82 (s, 3H), 2.45 (t, *J* = 7.6 Hz, 2H), 1.33 (s, 9H), 1.10 (t, J=7.6 Hz, 3H).

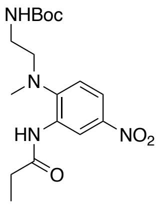

*tert-Butyl ethyl(2-(methyl(4-nitro-2-propionamidophenyl)amino)ethyl)carbamate (**38c**):*

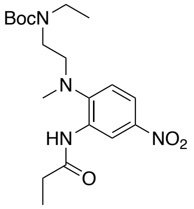

Intermediate **38c** was prepared from **37** and tert-butyl ethyl(2-(methylamino)ethyl)carbamate (**37d**) in the same manner as described for the preparation of **38a**. Yellow oil (456 mg, 18.39%). $^{1}$H NMR (400 MHz, DMSO-$d_6$) *δ* 9.27 (s, 1H), 8.53−8.38 (m, 1H), 7.99−7.84 (m, 1H), 7.26−7.08 (m, 1H), 3.28 (brs, 4H), 3.14−2.95 (m, 2H), 2.89 (s, 3H), 2.46−2.39 (m, 2H), 1.34 (s, 9H), 1.10 (t, *J* = 7.6 Hz, 3H), 0.96 (t, *J* = 7.2 Hz, 3H). **LCMS** $R_t$ = 1.022 min in 1.5 min chromatography, purity 71.2%, MS ESI *m/z*: 395.2 $[M+H]^+$.

*N-(2-(methyl(2-(pyrrolidin-1-yl)ethyl)amino)-5-nitrophenyl)propionamide (**38d**):* Intermediate **38d**

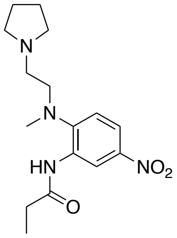

was prepared from **37** and *N*-methyl-2-(pyrrolidin-1-yl)ethan-1-amine in the same manner as described for the preparation of **38a**. Black brown oil (1.48 g, crude). $^{1}$H NMR (400 MHz, DMSO-$d_6$) *δ* 9.84 (s, 1H), 8.69 (br d, *J* = 2.4 Hz, 1H), 7.91 (dd, *J* = 2.8, 8.8 Hz, 1H), 7.18 (d, *J* = 9.2 Hz, 1H), 3.36 (br s, 2H), 3.18 (t, *J* = 6.8 Hz, 2H), 2.84 (s, 3H), 2.64 (t, *J* = 6.4 Hz, 2H), 2.48−2.46 (m, 2H),2.42−2.40 (m, 2H), 1.67−1.64 (m, 4H), 1.12-1.08 (m, 3H). LCMS $R_t$ = 1.069 min in 2.5 min chromatography, purity 51.5%, MS ESI *m/z*: 312.2 $[M+H]^+$.

*N-(2-(methyl(2-(piperidin-1-yl)ethyl)amino)-5-nitrophenyl)propionamide (**38e**):* Intermediate **38e**

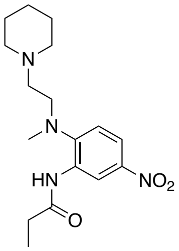

was prepared from **37** and *N*-methyl-2-(piperidin-1-yl)ethan-1-amine in the same manner as described for the preparation of **38a**. Brown solid (650 mg, 47.9%). $^{1}$H NMR (400 MHz, DMSO-$d_6$) *δ* 9.46 (s, 1H), 8.46 (d, *J* = 2.4 Hz, 1H), 7.93−7.90 (m, 1H), 7.14 (d, *J* = 9.2 Hz, 1H), 3.25 (t, *J* = 6.8 Hz, 2H), 2.86 (s, 3H), 2.45−2.40 (m, 4H), 2.35−2.24 (m, 4H), 1.40−1.33 (m, 6H), 1.10 (t, *J* = 7.6 Hz, 3H). LCMS $R_t$ = 1.350 min in 2.5 min chromatography, purity 18.7%, MS ESI *m/z*: 335.1 $[M+H]^+$.

*tert-Butyl (1-(4-nitro-2-propionamidophenyl)piperidin-3-yl)carbamate (**38f**):* Intermediate **38f** was

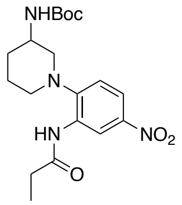

prepared from **37** and *tert*-butyl piperidin-3-ylcarbamate in the same manner as described for the preparation of **38a**. Black solid (2.2g, crude). $^{1}$H NMR (400 MHz, DMSO-$d_6$) *δ* 9.12 (s, 1H), 8.59 (d, *J* = 2.4 Hz, 1H), 7.95 (dd, *J* = 2.8, 8.8 Hz, 1H), 7.19 (d, *J* = 8.8 Hz, 1H), 7.07 (br d, *J* = 8.0 Hz, 1H), 3.84-3.69 (m, 1H), 3.21−3.13 (m, 1H), 3.03−2.97 (m, 1H), 2.89−2.67 (m, 2H), 2.49−2.39 (m, 2H), 1.86−1.63 (m, 4H), 1.38 (s, 9H), 1.11 (t, *J* = 7.6 Hz, 3H). LCMS $R_t$ = 0.567 min in 4 min chromatography, purity 74.2%, MS ESI *m/z*: 393.1 $[M+H]^+$.

*tert-Butyl 4-(4-nitro-2-propionamidophenyl)piperazine-1-carboxylate (**38g**):* Intermediate **38g** was prepared from **37** and *tert*-butyl piperazine-1-carboxylate in the same manner as described for the preparation of **38a**. Brown oil (1.7g, crude). $^{1}$H NMR (400 MHz, DMSO-$d_6$) $\delta$ 9.11 (s, 1H), 8.69 (d, $J$ = 2.4 Hz, 1H), 7.96 (dd, $J$ = 2.8, 8.8 Hz, 1H), 7.25 (d, $J$ = 9.2 Hz, 1H), 3.54 (br s, 4H), 2.99−2.90 (m, 4H), 2.46 (d, $J$ = 7.6 Hz, 2H), 1.38 (s, 9H), 1.11 (t, $J$ = 7.2 Hz, 3H). LCMS $R_t$ = 0.561 min in 4 min chromatography, purity 99.8%, MS ESI $m/z$: 379.1 $[M+H]^+$.

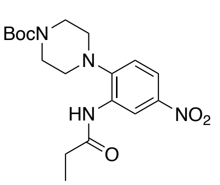

*tert-Butyl methyl(1-(4-nitro-2-propionamidophenyl)piperidin-3-yl)carbamate (**38h**):* Intermediate **38h** was prepared from **37** and *tert*-butyl methyl(piperidin-3-yl)carbamate in the same manner as described for the preparation of **38a**. Brown solid (2.90 g, crude).$^{1}$H NMR (400 MHz, DMSO-$d_6$) $\delta$ 9.09 (br s, 1H), 8.61 (d, $J$ = 2.4 Hz, 1H), 7.95 (dd, $J$ = 2.8, 8.8 Hz, 1H), 7.25 (d, $J$ = 8.8 Hz, 1H), 4.36−4.23 (m, 1H), 3.783.68 (m, 1H), 3.20−3.07 (m, 2H), 2.77−2.63 (m, 2H), 2.47−2.30 (m, 2H), 1.81−1.74 (m, 2H), 1.73−1.60 (m, 2H), 1.40 (s, 9H), 1.18 (t, $J$ = 7.2 Hz, 3H). LCMS $R_t$ = 0.604 min in 1 min chromatography, purity 63.7%, MS ESI $m/z$: 407.2 $[M+H]^+$.

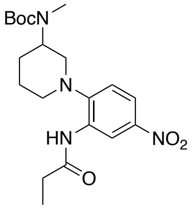

*tert-Butyl (2-((4-amino-2-propionamidophenyl)(methyl)amino)ethyl)(methyl)carbamate (**39a:**)* To a solution of compound **38a** (700 mg, 1.84 mmol) in MeOH (3 mL) was added Pd/C (630 mg, 10% purity) under $H_2$ atmosphere. The suspension was degassed and purged with $H_2$ for 3 hours. Then the reaction mixture was stirred under $H_2$ (15 Psi) at 25℃ for 10 hours. The reaction mixture was filtered through a pad of Celite®, washed with MeOH (20 mL), and the combined organic extracts evaporated to dryness in *vacuo* to afford the crude product **39a** (579 mg, 97.2%) as a gray solid. $^{1}$H NMR (400 MHz, DMSO-$d_6$) $\delta$ 8.69 (br s, 1H), 7.53 (s, 1H), 6.95 (d, $J$ = 8.4 Hz, 1H), 6.23 (dd, $J$ = 2.4, 8.4 Hz, 1H), 4.90 (s, 2H), 3.25−3.19 (m, 2H), 2.82 (br t, $J$ = 6.4 Hz, 2H), 2.76 (s, 3H), 2.51 (br s, 3H), 2.35 (br d, $J$ = 7.2 Hz, 2H), 1.35 (br s, 9H), 1.08 (t, $J$ = 7.6 Hz, 3H). LCMS $R_t$ = 0.427 min in 1 min chromatography, purity 97.21%, MS ESI $m/z$: 235.1 $[M+H]^+$.

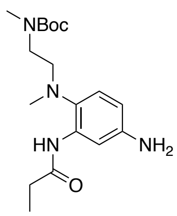

*tert-Butyl (2-((4-amino-2-propionamidophenyl)(methyl)amino)ethyl)carbamate (**39b**):* Intermediate **39b** was prepared from **38b** in the same manner as described for the preparation of **39a**. Brown solid (1.2 g, crude). $^{1}$H NMR (400 MHz, DMSO-$d_6$) δ 8.86 (s, 1H), 7.51 (d, $J$ = 1.6 Hz, 1H), 6.93−6.87 (m, 2H), 6.23 (dd, $J$ = 2.4, 8.4 Hz, 1H), 4.87 (s, 2H), 2.99−2.96 (m, 2H), 2.73 −2.70 (m, 2H), 2.48 (s, 3H), 2.39 (q, $J$ = 7.6 Hz, 2H), 1.36 (s, 9H), 1.09 (t, $J$ = 7.6 Hz, 3H). LCMS $R_t$ = 0.413 min in 1 min chromatography, purity 96.5%, MS ESI $m/z$: 337.1 $[M+H]^+$.

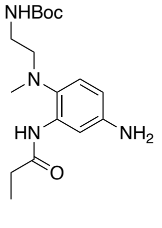

*tert-Butyl (2-((4-amino-2-propionamidophenyl)(methyl)amino)ethyl)(ethyl)carbamate (**39c**):*

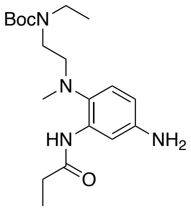

Intermediate **39c** was prepared from **38c** in the same manner as described for the preparation of **39a**. Purple oil (456 mg, 18.39%). **$^1$H NMR** (400 MHz, DMSO-$d_6$) $\delta$ 8.73 (br s, 1H), 7.51 (s, 1H), 7.05-6.91 (m, 1H), 6.26−6.18 (m, 1H), 4.92 (brs, 2H), 3.20−3.11 (m, 4H), 2.85−2.77 (m, 2H), 2.53−2.52 (m, 2H), 2.42−2.25 (m, 3H), 1.39−1.31 (m, 9H), 1.08 (t, $J$ = 7.2 Hz, 3H), 0.99 (t, $J$ = 7.2 Hz, 3H). LCMS $R_t$ = 1.137 min in 2.5 min chromatography, purity 85.8%, MS ESI *m/z*: 365.2 $[M+H]^+$.

*N-(5-amino-2-(methyl(2-(pyrrolidin-1-yl)ethyl)amino)phenyl)propionamide (**39d**):* Intermediate

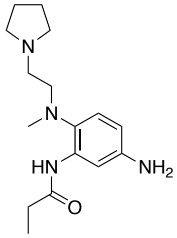

**39d** was prepared from **38d** in the same manner as described for the preparation of **39a**. Brown solid (1.48 g, crude). $^1$H NMR (400 MHz, DMSO-$d_6$) $\delta$ 9.46 (s, 1H), 7.50 (d, $J$ = 2.0 Hz, 1H), 6.96 (d, $J$ = 8.8 Hz, 1H), 6.24 (dd, $J$ = 2.4, 8.4 Hz), 4.92−4.90 (m, 2H), 2.81−2.75 (m, 2H), 2.56−2.52 (m, 3H), 2.43−2.41 (m, 4H), 2.31−2.27 (m, 4H), 1.75−1.62 (m, 4H), 1.09 (t, $J$ = 7.6 Hz, 3H).

*N-(5-amino-2-(methyl(2-(piperidin-1-yl)ethyl)amino)phenyl)propionamide (**39e**):* Intermediate

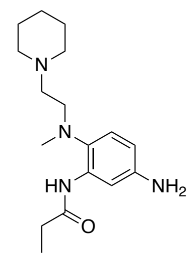

**39e** was prepared from **38e** in the same manner as described for the preparation of **39a**. Brown solid (276 mg, 89.0%). $^1$H NMR (400 MHz, DMSO-$d_6$) $\delta$ 9.08 (s, 1H), 7.46 (s, 1H), 6.94 (d, $J$ = 8.0 Hz, 1H), 6.24 (dd, $J$ = 2.4, 8.4 Hz, 1H), 4.92−4.90 (m, 2H), 2.83−2.79 (m, 2H), 2.53−2.51 (m, 3H), 2. 35−2.32 (m, 3H), 2.26 (s, 3H), 2.18−2.15 (m, 2H), 1.48−1.44 (m, 4H), 1.36 (br d, $J$ = 4.8 Hz, 2H), 1.12−1.08 (m, 3H). LCMS $R_t$ = 0.328 min in 1 min chromatography, purity 82.0%, MS ESI *m/z*: 305.2 $[M+H]^+$.

*tert-Butyl (1-(4-amino-2-propionamidophenyl)piperidin-3-yl)carbamate (**39f**):* Intermediate **39f**

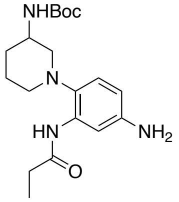

was prepared from **38f** in the same manner as described for the preparation of **39a**. Light brown solid (900 mg, 44.0%). $^1$H NMR (400 MHz, DMSO-$d_6$) $\delta$ 8.74 (br s, 1H), 7.37 (br s, 1H), 6.99 (br d, $J$ = 6.8 Hz, 1H), 6.80 (d, $J$ = 8.4 Hz, 1H), 6.22 (dd, $J$ = 2.4, 8.4 Hz, 1H), 4.86 (s, 2H), 3.65−3.60 (m, 1H), 2.81−2.70 (m, 1H), 2.67−2.61 (m, 1H), 2.53−2.50 (m, 1H), 2.46−2.28 (m, 3H), 1.84 (m, 3H), 1.37 (s, 9H), 1.35−1.27 (m, 1H), 1.11 (t, $J$ = 7.6 Hz, 3H). LCMS $R_t$ = 0.440 min in 1 min chromatography, purity 99.3%, MS ESI *m/z*: 363.1 $[M+H]^+$.

*tert-Butyl 4-(4-amino-2-propionamidophenyl)piperazine-1-carboxylate (**39g**):* To a solution of

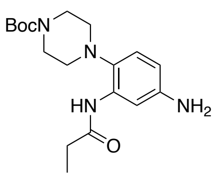

compound **38g** (1.10 g, 2.91 mmol) in EtOH (8.00 mL) was added $NH_4Cl$ (466 mg, 8.72 mmol) dissolved in $H_2O$ (2 mL) and Fe (974 mg, 17.44 mmol) at 25℃. Then the mixture was stirred at 80°C under $N_2$ atmosphere for 10 hours. The mixture cooled to rt and was filtered and the solid was washed with MeOH (15 mL×2). The residue was purified by prep-HPLC (column: Xtimate C18 150*40mm*10um; mobile phase: [water ($NH_3H_2O$)-ACN]; B%: 35%-65%, 10min) to obtain **39g** (500 mg, 48.5%) as a gray solid. $^1$H NMR (400 MHz, DMSO-$d_6$) $\delta$ 8.73 (s, 1H), 7.44 (br s, 1H), 6.88 (d, $J$ = 8.4 Hz, 1H), 6.23 (dd, $J$ = 2.4, 8.4 Hz, 1H), 4.95 (br s, 2H), 3.46 (br s, 4H), 2.62 (br t, $J$ = 4.8 Hz, 4H), 2.38 (q, $J$ = 7.6 Hz, 2H), 1.42 (s, 9H), 1.10 (t, $J$ = 7.6 Hz, 3H). LCMS $R_t$ = 0.451 min in 1 min chromatography, purity 98.2%, MS ESI *m/z*: 349.1 $[M+H]^+$.

*tert-Butyl (1-(4-amino-2-propionamidophenyl)piperidin-3-yl)(methyl)carbamate (**39h**):*

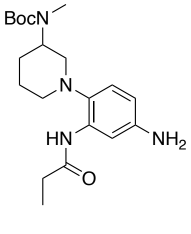

Intermediate **39h** was prepared from **38h** in the same manner as described for the preparation of **39g**. Brown solid (884 mg, 48.3%). $^1$H NMR (400 MHz, DMSO-$d_6$) $\delta$ 8.87−8.58 (m, 1H), 7.41 (br s, 1H), 6.88 (br d, $J$ = 8.0 Hz, 1H), 6.23 (dd, $J$ = 2.4, 8.4 Hz, 1H), 4.93 (s, 2H), 4.18−3.96 (m, 1H), 2.71 (s, 3H), 2.69−2.59 (m, 2H), 2.49−2.35(m, 2H), 1.79−1.76 (m, 3H), 1.71−1.68 (m, 1H), 1.59 (s, 9H), 1.13 (t, $J$ = 7.2 Hz, 3H). LCMS $R_t$ = 0.458 min in 1 min chromatography, purity 86.2%, MS ESI *m/z*: 377.1 $[M+H]^+$.

*5-Chloro-7-(cyclopropylamino)pyrazolo[1,5-a]pyrimidine-3-carbonitrile (**41**):* To a solution of

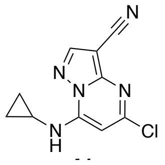

compound **40** (5.00 g, 23.47 mmol, 1 *eq*) in EtOH (30 mL) was added compound cyclopropylamine (12.00 g, 211.24 mmol, 14 mL, 9 *eq*) dropwise at 25℃. Then the mixture was stirred at 25°C for 2 hours. The reaction mixture was filtered, and the solid was washed with EtOH (4 mL × 2). The mixture was filtered, the solids washed with EtOAc, and dried in vacuo to afford the title compound (crude) as a yellow solid (5.47 g, 23.03 mmol, 98.14% yield) which was used without further purification. $^1$H NMR (400 MHz, MeOD-$d_4$) $\delta$ 8.36 (s, 1H), 6.61 (s, 1H), 2.76 (m, 1H), 1.04−0.93 (m, 2H), 0.83−0.74 (m, 2H). LCMS $R_t$ = 0.500 min in 1 min chromatography, Chromolith @ Flash RP-18e, 25-3mm, purity 97.93%, MS ESI *m/z*: 234.0 $[M+H]^+$.

*tert-Butyl (2-((4-((3-cyano-7-(cyclopropylamino)pyrazolo[1,5-a]pyrimidin-5-yl)amino)-2-propionamidophenyl)(methyl)amino)ethyl)(methyl)carbamate (**42a**):* To a solution of compound

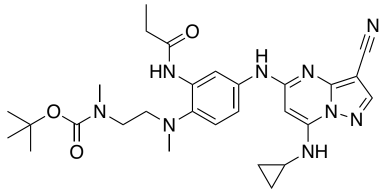

**39a** (250 mg, 1.07 mmol) and compound **40** (562 mg, 1.60 mmol) in dioxane (3 mL) was added $Cs_2CO_3$ (1.05 g, 3.21 mmol), BINAP (100 mg, 0.16 mmol) and $Pd(OAc)_2$ (36 mg, 0.16 mmol) at 25℃. Then the mixture was degassed and purged with $N_2$. The reaction mixture was heated in a microwave at 130°C for 0.5h. The mixture was cooled to rt and water (10 mL) was added to the reaction, which was extracted with DCM (4×15 mL). The organic layer was washed with water (2×10 mL), dried with $Na_2SO_4$, filtered, and concentrated to give crude product, which was purified by flash chromatography (eluent of 0~5%, MeOH/DCM) to give the desired product **42a** (757 mg, 49.0%) as a yellow solid. $^1$H NMR (400 MHz, DMSO-$d_6$) $\delta$ 9.66 (s, 1H), 8.82 (br s, 1H), 8.33 (s, 1H), 8.18 (d, $J$ = 0.8 Hz, 2H), 7.89 (br d, $J$ = 4.8 Hz, 1H), 7.25 (br d, $J$ = 8.0 Hz, 1H), 6.05 (s, 1H), 4.94 (br s, 1H), 3.56 (s, 2H), 3.33 (m, 3H), 2.62 (s, 3H), 2.147−2.39 (m, 2H), 1.35 (br s, 9H), 1.131.06 (m, 3H), 0.81−0.79 (m, 2H), 0.73-0.71 (m, 2H). LCMS $R_t$ = 1.498 min in 4 min chromatography, purity 28%, MS ESI *m/z*: 548.3 $[M+H]^+$.

*tert-Butyl (2-((4-((3-cyano-7-(cyclopropylamino)pyrazolo[1,5-a]pyrimidin-5-yl)amino)-2-propionamidophenyl)(methyl)amino)ethyl)carbamate (**42b**):* Intermediate **42b** was prepared

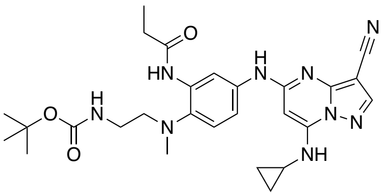

from **39b** and **41** in the same manner as described for the preparation of **42a**. Brown solid (212 mg, 32.8%). $^1$H NMR (400 MHz, DMSO-$d_6$) $\delta$ 9.64 (s, 1H), 8.96 (s, 1H), 8.33 (s, 1H), 8.18 (s, 2H), 7.89−7.78 (m, 1H), 7.21 (d, $J$ = 8.8 Hz, 1H), 6.99 - 6.91 (m, 1H), 6.04 (s, 1H), 3.32−3.30 (m, 1H), 3.10-3.03 (m, 2H), 2.85−2.79 (m, 2H), 2.58 (s, 3H), 2.48−2.43 (m, 2H), 1.36 (s, 9H), 1.12 (t, $J$ = 7.6 Hz, 3H), 0.83−0.77 (m, 2H), 0.73−0.68 (m, 2H). LCMS $R_t$ = 2.361 min in 4 min chromatography, purity 70.8%, MS ESI *m/z*: 534.3 $[M+H]^+$.

*tert-Butyl (2-((4-((3-cyano-7-(cyclopropylamino)pyrazolo[1,5-a]pyrimidin-5-yl)amino)-2-propionamidophenyl)(methyl)amino)ethyl)(ethyl)carbamate (**42c**):* Intermediate **42c** was

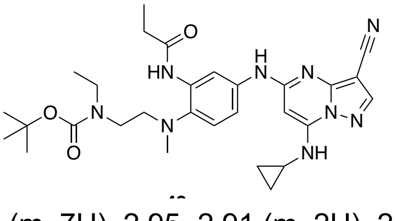

prepared from **39c** in the same manner as described for the preparation of **42a**. Purple oil (456 mg, 18.39%). $^1$H NMR (400 MHz, DMSO-$d_6$) $\delta$ 9.64 (s, 1H), 8.85 (s, 1H), 8.33 (s, 1H), 8.19−8.14 (m, 2H), 7.29−7.18 (m, 2H), 6.03 (s, 1H), 3.21−3.10 (m, 7H), 2.95−2.91 (m, 2H), 2.62 (s, 3H), 1.37−1.34 (m, 9H), 1.14−1.10 (m, 3H), 1.03−1.00 (m,

3H), 0.82−0.78 (m, 2H), 0.72−0.69 (m, 2H). LCMS $R_t$ = 1.027 min in 1.5 min chromatography, purity 71.0%, MS ESI *m/z*: 562.4 $[M+H]^+$.

*tert-Butyl (1-(4-((3-cyano-7-(cyclopropylamino)pyrazolo[1,5-a]pyrimidin-5-yl)amino)-2-propionamidophenyl)piperidin-3-yl)carbamate (**42f**):* Intermediate **42f** was prepared from **39f** in

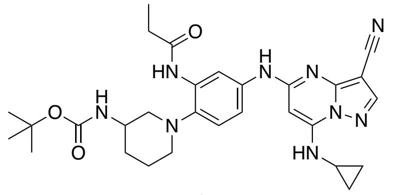

the same manner as described for the preparation of **42a**. Yellow solid I (1.27 g, 67.7%). [1]H NMR (400 MHz, DMSO-$d_6$) δ 9.62 (s, 1H), 8.88 (s, 1H), 8.33 (s, 1H), 8.18 (s, 1H), 8.02 (d, *J* = 2.0 Hz, 1H), 7.08 (d, *J* = 8.4 Hz, 2H), 6.02 (s, 1H), 5.76 (s, 1H), 3.75−3.72 (m, 1H), 2.91−2.78 (m, 1H), 2.78−2.73 (m, 1H), 2.66−2.58 (m, 2H), 2.47−2.40 (m, 3H), 1.81−1.68 (m, 4H), 1.38 (s, 9H), 1.13 (t, *J* = 7.6 Hz, 3H), 0.83−0.75 (m, 2H), 0.73−0.69 (m, 2H). LCMS $R_t$ = 2.481 min in 4 min chromatography, purity 51.0 %, MS ESI *m/z*: 560.4 $[M+H]^+$.

*tert-Butyl 4-(4-((3-cyano-7-(cyclopropylamino)pyrazolo[1,5-a]pyrimidin-5-yl)amino)-2-propionamidophenyl)piperazine-1-carboxylate (**42g**):* Intermediate **42g** was prepared from **39g**

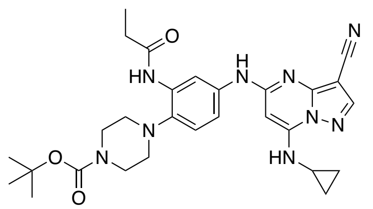

in the same manner as described for the preparation of **42a**. Brown solid (800 mg, 83.5%). [1]H NMR (400 MHz, DMSO-$d_6$) *δ* 9.41 (s, 1H), 8.60 (s, 1H), 8.09 (s, 1H), 7.94 (s, 1H), 7.86 (s, 1H), 7.65-7.57 (m, 1H), 6.92 (d, *J* = 8.4 Hz, 1H), 5.78 (s, 1H), 3.35−3.21 (m, 4H), 3.13−3.07 (m, 4H), 2.23−2.15 (m, 2H), 1.18 (s, 10H), 0.91−0.82 (m, 3H), 0.57−0.53 (m, 2H), 0.51−0.43 (m, 2H). LCMS $R_t$ = 2.468 min in 4 min chromatography, purity 60.9%, MS ESI *m/z*: 546.3 $[M+H]^+$ .

*tert-Butyl (1-(4-((3-cyano-7-(cyclopropylamino)pyrazolo[1,5-a]pyrimidin-5-yl)amino)-2-propionamidophenyl)piperidin-3-yl)(methyl)carbamate (**42h**):* Intermediate **42h** was prepared

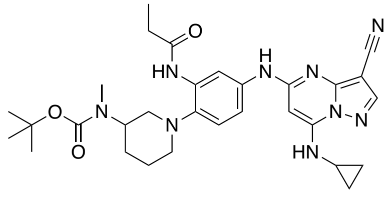

from **39h** in the same manner as described for the preparation of **42a**. Black brown solid (516 mg, 72.6%). LCMS $R_t$ = 4.568 min in 7 min chromatography, purity 69.1%, MS ESI *m/z*: 574.4 $[M+H]^+$.

*5-Chloro-7-(cyclobutylamino)pyrazolo[1,5-a]pyrimidine-3-carbonitrile (**44a**):* To a solution of

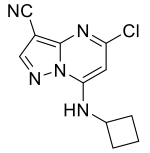

compound **43** (350 mg, 1.64 mmol) in EtOH (5 mL) was added cyclobutylamine (234 mg, 3.29 mmol) dropwise at 25℃ and then the resulting mixture was stirred for 2 hours. Water (10 mL) was added to the reaction, which was extracted with DCM (2×20 mL). The organic layer was washed with water (2×10 mL), dried with $Na_2SO_4$, filtered, and concentrated to give the crude product, which was purified by flash chromatography (eluent of 0~5%, MeOH/DCM) to give the desired product **44a** (314 mg,) as a light-yellow solid.

$^1$H NMR (DMSO-$d_6$, 400 MHz) $\delta$ 9.14 (d, $J$ = 6.0 Hz, 1H), 8.67 (s, 1H), 6.50 (s, 1H), 4.39−4.17 (m, 1H), 2.40−2.18 (m, 4H), 1.79− 1.57 (m, 2H). LCMS $R_t$ = 0.519 min in 1 min chromatography, Chromolith Flash RP-18, 5um, 3.0*25mm, purity 98.30 %, MS ESI *m/z*: 248.0 $[M+H]^+$

*5-Chloro-7-((3,3-difluorocyclobutyl)amino)pyrazolo[1,5-a]pyrimidine-3-carbonitrile (**44b**):*

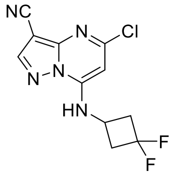

Intermediate **44b** was prepared from **43** and 3,3-difluorocyclobutan-1-amine in the same manner as described for the preparation of **44a**. Light yellow solid. (158 mg, 0.55 mmol, 58.49% yield, 98.58% purity). $^1$H NMR (400 MHz, DMSO-$d_6$) $\delta$ 9.33 (br s, 1H), 8.71 (s, 1H), 6.66 (s, 1H), 4.35−4.16 (m, 1H), 3.20−2.89 (m, 4H). LCMS $R_t$ = 0.511 min in 1 min chromatography, Chromolith Flash RP-18, 5um, 3.0*25mm, purity 97.56%, MS ESI *m/z*: 284.0 $[M+H]^+$

*7-(Bicyclo[1.1.1]pentan-1-ylamino)-5-chloropyrazolo[1,5-a]pyrimidine-3-carbonitrile (**44c**):*

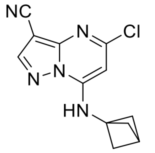

Intermediate **44c** was prepared from **43** and bicyclo[1.1.1]pentan-1-amine in the same manner as described for the preparation of **44a**. White solid (216 mg, 0.82 mmol, 87.69% yield, 98.98% purity). $^1$H NMR (400 MHz, DMSO-$d_6$) $\delta$ 9.57 (br s, 1H), 8.69 (s, 1H), 6.52 (s, 1H), 2.23 (s, 7H). LCMS $R_t$ = 0.537 min in 1 min chromatography, Chromolith Flash RP-18, 5um, 3.0*25mm, purity 98.582%, MS ESI *m/z*: 260.1 $[M+H]^+$.

*5-Chloro-7-((1-methylcyclopropyl)amino)pyrazolo[1,5-a]pyrimidine-3-carbonitrile (**44d**):*

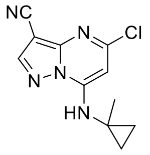

Intermediate **44d** was prepared from **43** and 1-methylcyclopropan-1-amine in the same manner as described for the preparation of **44a**. Yellow solid (340 mg, 0.8 mmol, 55.96% yield, 57.40% purity). **$^1$H** NMR (DMSO-$d_6$, 400 MHz) $\delta$ 8.20−7.80 (m, 1H), 5.70−5.57 (m, 1H), 1.22 (s, 3H), 0.56−0.47 (m, 2H), 0.46−0.38 (m, 2H). LCMS $R_t$ = 0.506 min in 1 min chromatography, Chromolith Flash RP-18, 5um, 3.0*25mm, purity 57.40%, MS ESI *m/z*: 248.0 $[M+H]^+$.

*7-(((3s,5s,7s)-Adamantan-1-yl)amino)-5-chloropyrazolo[1,5-a]pyrimidine-3-carbonitrile (**44e**):*

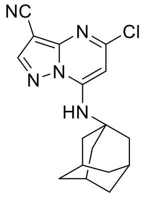

Intermediate **44e** was prepared from **43** and (1R,5S)-bicyclo[3.3.1]nonan-3-amine in the same manner as described for the preparation of **44a**. White solid (697 mg, crude). $^1$H NMR (DMSO-$d_6$, 400 MHz) $\delta$ 8.85−8.54 (m, 1H), 7.35−7.00 (m, 1H), 6.76 (s, 1H), 4.70−4.51 (m, 1H), 4.03 (d, $J$ = 7.2 Hz, 1H), 2.15−2.07 (m, 4H), 1.98 (s, 2H), 1.84−1.71 (m, 2H), 1.70−1.58 (m, 2H), 1.51−1.43 (m, 1H), 1.25−1.13 (m, 2H). LCMS $R_t$ = 0.625 min in 1 min chromatography, Chromolith Flash RP-18, 5um, 3.0*25mm, purity 78.13 %, MS ESI *m/z*: 328.1 $[M+H]^+$.

*7-(((1r,3r,5r,7r)-Adamantan-2-yl)amino)-5-chloropyrazolo[1,5-a]pyrimidine-3-carbonitrile (**44f**):*

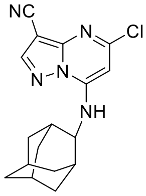

Intermediate **44f** was prepared from **43** and (1R,5S)-bicyclo[3.3.1]nonan-3-amine in the same manner as described for the preparation of **44a**. Light yellow solid (317 mg, crude). $^1$H NMR (DMSO-$d_6$, 400 MHz) $\delta$ 8.72 (s, 1H), 6.73 (s, 1H), 4.08−4.04 (m, 1H), 2.90−2.81 (m, 2H), 2.13−2.04 (m, 2H), 1.98−1.92 (m, 5H), 1.88−1.85 (m, 2H), 1.76−1.69 (m, 2H), 1.64−1.57 (m, 2H). LCMS $R_t$ = 0.630 min in 1 min chromatography, Chromolith Flash RP-18, 5um, 3.0*25mm, purity 97.71 %, MS ESI *m/z*: 328.0 $[M+H]^+$.

*N-(1-(3-cyano-7-(phenylamino)pyrazolo[1,5-a]pyrimidin-5-yl)-1H-indol-6-yl)acetamide (**44g**):*

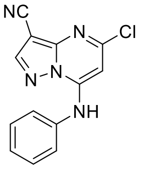

Intermediate **44g** was prepared from **43** and aniline in the same manner as described for the preparation of **44a**. Light yellow solid (316 mg, 1.15 mmol, 94.46% yield, 98.39% purity). $^1$H NMR (400 MHz, DMSO-$d_6$) $\delta$ 10.88 (br s, 1H), 8.80 (s, 1H), 7.57−7.43 (m, 4H), 7.42−7.30 (m, 1H), 6.29 (s, 1H). LCMS $R_t$ = 0.523 min in 1 min chromatography, Chromolith Flash RP-18, 5um,3.0*25mm, purity 98.39%, MS ESI *m/z*: 270.0 $[M+H]^+$.

*5-Chloro-7-((oxetan-3-ylmethyl)amino)pyrazolo[1,5-a]pyrimidine-3-carbonitrile (**44h**):*

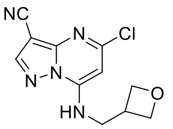

Intermediate **44h** was prepared from **43** and oxetan-3-ylmethanamine in the same manner as described for the preparation of **44a**. Light yellow solid (300 mg, 0.90 mmol, 87.16% yield, 95.64% purity). $^1$H NMR (400 MHz, DMSO-$d_6$) $\delta$ 8.66 (s, 1H), 6.71 (br s, 1H), 4.63 (t, $J$ = 6.8 Hz, 2H), 4.35 (t, $J$ = 6.0 Hz, 2H), 3.73 (br d, $J$ = 7.6 Hz, 2H), 3.29 (m, 1H). LCMS $R_t$ = 0.450 min in 1 min chromatography, Chromolith Flash RP-18, 5um, 3.0*25mm, purity 53.02%, MS ESI *m/z*: 264.0 $[M+H]^+$.

*5-Chloro-7-(((tetrahydrofuran-3-yl)methyl)amino)pyrazolo[1,5-a]pyrimidine-3-carbonitrile (**44i**):*

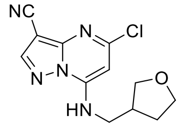

Intermediate **44i** was prepared from **43** and (tetrahydrofuran-3-yl)methanamine in the same manner as described for the preparation of **44a**.Yellow (830 mg, crude). $^1$H NMR (DMSO-$d_6$, 400 MHz) $\delta$ 9.04 (s, 1H), 8.66 (s, 1H), 6.69 (s, 1H), 3.823.72 (m, 1H), 3.72 −3.65 (m, 1H), 3.63−3.59 (m, 1H), 3.52−3.43 (m, 3H), 2.70−2.56 (m, 1H), 2.14−1.86 (m, 1H), 1.73−1.52 (m, 1H). LCMS $R_t$ = 0.466 min in 1 min chromatography, Chromolith Flash RP-18, 5um, 3.0*25mm, purity 98.07 %, MS ESI *m/z*: 278.0 $[M+H]^+$.

*5-Chloro-7-(((tetrahydro-2H-pyran-4-yl)methyl)amino)pyrazolo[1,5-a]pyrimidine-3-carbonitrile*

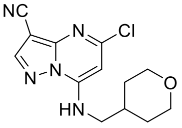

*(**44j**):* Intermediate **44j** was prepared from **43** and (tetrahydro-2H-pyran-4-yl)methanamine in the same manner as described for the preparation of **44a**.White solid (273 mg, crude). $^1$H NMR (DMSO-$d_6$, 400 MHz) $\delta$ 9.05 - 8.88

(m, 1H), 8.65 (s, 1H), 6.68 (s, 1H), 3.93−3.76 (m, 2H), 3.35−3.31 (m, 2H), 3.27−3.19 (m, 2H), 2.02−1.86 (m, 1H), 1.70−1.52 (m, 2H), 1.32−1.13 (m, 2H). LCMS $R_t$ = 0.484 min in 1 min chromatography, Chromolith Flash RP-18, 5um, 3.0*25mm, purity 99.27 %, MS ESI *m/z*: 292.0 $[M+H]^+$.

*5-Chloro-7-((1-methylpiperidin-4-yl)amino)pyrazolo[1,5-a]pyrimidine-3-carbonitrile (**44k**):*

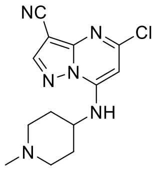


Intermediate **44k** was prepared from **43** and 1-methylpiperidin-4-amine in the same manner as described for the preparation of **44a**. White solid (672 mg, 2.13 mmol, 64.93% yield, 92.31% purity). $^1$H NMR (400 MHz, DMSO-$d_6$) $\delta$ 8.67 (br s, 1H), 8.64 (s, 1H), 6.70 (s, 1H), 3.68−3.62 (m, 1H), 2.78−2.70 (m, 2H), 2.14 (s, 3H), 2.05−1.93 (m, 2H), 1.85−1.71 (m, 4H). LCMS $R_t$ = 0.372 min in 1 min chromatography, Chromolith Flash RP-18, 5um, 3.0*25mm, purity 92.31%, MS ESI *m/z*: 291.1 $[M+H]^+$.

*7-((1-Acetylpiperidin-4-yl)amino)-5-chloropyrazolo[1,5-a]pyrimidine-3-carbonitrile (**44l**):*

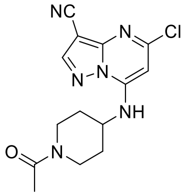


Intermediate **44l** was prepared from **43** and 1-(4-aminopiperidin-1-yl)ethan-1-one in the same manner as described for the preparation of **44a**. Light yellow solid (300 mg, 0.90 mmol, 87.16% yield, 95.64% purity). $^1$H NMR (400 MHz, DMSO-$d_6$) $\delta$ 8.66 (s, 1H), 8.57−7.78 (m, 1H), 6.79 (s, 1H), 4.47−4.41 (m, 1H), 3.98 (br t, $J$ = 10.8 Hz, 1H), 3.87 (br d, $J$ = 13.2 Hz, 1H), 3.15 (t, $J$ = 12.8 Hz, 1H), 2.63 (t, $J$ = 12.4 Hz, 1H), 2.01 (s, 3H), 1.94−1.80 (m, 2H), 1.75−1.49 (m, 2H). LCMS $R_t$ = 0.447 min in 1 min chromatography, Chromolith Flash RP-18, 5um, 3.0*25mm, purity 95.64%, MS ESI *m/z*: 391.1 $[M+H]^+$.

*5-Chloro-7-(((1-methyl-1H-pyrazol-4-yl)methyl)amino)pyrazolo[1,5-a]pyrimidine-3-carbonitrile (**44m**):*

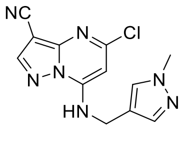


Intermediate **44m** was prepared from **43** and (1-methyl-1H-pyrazol-4-yl)methanamine in the same manner as described for the preparation of **44a**. Light yellow solid (293 mg, crude). $^1$H NMR (DMSO-$d_6$, 400 MHz) $\delta$ 8.66 (s, 1H), 7.81−7.63 (m, 1H), 7.55−7.39 (m, 1H), 6.64 (s, 1H), 4.49 (d, $J$ = 6.4 Hz, 2H), 3.82 (s, 1H), 3.77 (s, 3H). LCMS $R_t$ = 0.447 min in 1 min chromatography, Chromolith Flash RP-18, 5um, 3.0*25mm, purity 89.51 %, MS ESI *m/z*: 288.0 $[M+H]^+$.

*Ethyl (5-chloro-3-cyanopyrazolo[1,5-a]pyrimidin-7-yl)-D-alaninate (**44n**):* Intermediate **44n** was

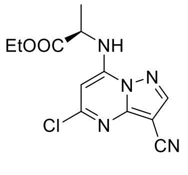


prepared from **43** and ethyl D-alanine in the same manner as described for the preparation of **44a**. Light yellow solid (882 mg, 2.89 mmol, 61.47% yield, 96.08% purity). LCMS $R_t$ = 0.507 min in 1 min chromatography, purity 97.37%, MS ESI *m/z*: 239.9 $[M+H]^+$.

*Ethyl (5-(6-acetamido-1H-indol-1-yl)-3-cyanopyrazolo[1,5-a]pyrimidin-7-yl)-D-alaninate (**47**):* To

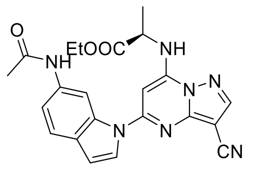

a solution of **44n** (620 mg, 2.11 mmol, 1 *eq*) and **46** (367 mg, 2.11 mmol, 1 *eq*) in dioxane (8 mL) was added $Cs_2CO_3$ (2.06 g, 6.33 mmol, 3 *eq*) Xantphos (183 mg, 0.31 mmol, 0.15 *eq*) and $Pd(OAc)_2$ (71 mg, 0.31 mmol, 0.15 *eq*) at 25°C. The mixture was purged with $N_2$ and heated in a microwave condition at 130°C for 0.5 hr. The mixture was cooled to rt and water (10 mL) was added to the reaction, which was extracted with DCM (3×15 mL). The organic layer was washed with water (2×10 mL), dried with $Na_2SO_4$, filtered, and concentrated to give crude product, which was purified by flash silica gel chromatography (eluent: 0-7%, MeOH / DCM) to get **47** (804 mg, crude purity) as a yellow solid. $^1$H NMR (400 MHz, DMSO-$d_6$) $\delta$ 9.98 (s, 1H), 8.85−8.83 (m, 1H), 8.61−8.59 (m, 1H), 8.05−8.03 (m, 1H), 7.54 (d, *J* = 8.4 Hz, 1H), 7.32−7.27 (m, 2H), 6.89−6.87 (m, 1H), 6.87−6.76 (m, 1H), 4.98−4.95 (m, 1H), 4.23 - 3.33 (m, 2H), 2.07−2.04 (m, 3H), 1.60 (d, *J* = 7.2 Hz, 3H), 1.33−1.15 (m, 3H).

*N-(1H-indol-6-yl)acetamide (**46**):* To a solution of 1H-indol-6-amine **45** (5.00 g, 37.83 mmol) in

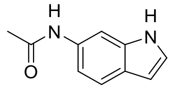

toluene (60 mL) was added acetyl acetate (7.72 g, 75.66 mmol, 7.09 mL). The reaction mixture was stirred at 25°C under $N_2$ atmosphere. After 2 h, the reaction mixture was partitioned between water (20 mL) and EtOAc (100 mL). The layers were separated, and the aqueous phase was further extracted with EtOAc (2×100 mL). The combined organic phases were washed twice with water and brine (20 mLx2), $Na_2SO_4$, filtered, and concentrated. The residue was purified by flash silica gel chromatography (eluent of 0~9% MeOH in DCM) to get the purified compound. After concentration, the compound was recrystallized from acetonitrile to afford **46** (5.6 g, 32.11 mmol, 84.87% yield, 99.88 % purity) as a grey solid. $^1$H NMR (DMSO-$d_6$, 400 MHz) $\delta$ 10.95 (s, 1H), 9.80 (s, 1H), 7.97 (s, 1H), 7.41 (d, *J* = 8.4 Hz, 1H), 7.23 (t, *J* = 2.8 Hz, 1H), 6.99 (dd, *J* = 1.6, 8.4 Hz, 1H), 6.33 (t, *J* = 2.0 Hz, 1H), 2.04 (s, 3H). LCMS $R_t$ = 0.981 min in 4 min chromatography, Xtimate C18, 3um,2.1*30mm, purity 99.87%, MS ESI *m/z*: 175.3 $[M+H]^+$. HPLC $R_t$ = 1.691 min in 8 min chromatography, XBridge Shield RP18 (2.1×50mm 5µm), purity 99.64%.

*(R)-2-((5-(6-Acetamido-1H-indol-1-yl)-3-cyanopyrazolo[1,5-a]pyrimidin-7-yl)amino)propenamide*

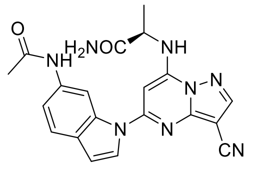

*(**48**):* A mixture of compound **47** (750 mg, 1.74 mmol, 1 eq) in $NH_3$/MeOH (7 M, 5 mL) was stirred at 25°C for 1hr. The reaction mixture was filtered by filter paper. The filter cake was the target coarse product which was washed with MeOH (3×5 mL) and concentrated directly. The crude was used in the next step directly. Compound **48** (390 mg, 0.90 mmol, 49.36% yield, 88.53% purity)

was obtained as a brown solid. $^{1}$H NMR (DMSO-$d_6$, 400 MHz) $\delta$ 9.98 (s, 1H), 8.82 (s, 1H), 8.68 (s, 1H), 8.03 (d, $J$ = 3.6 Hz, 1H), 7.68−7.63 (m, 2H), 7.48−7.42 (m, 1H), 6.91−6.84 (m, 2H), 6.78 (d, $J$ = 3.6 Hz, 1H), 6.58 (s, 1H), 4.65−4.46 (m, 1H), 2.08 (s, 3H), 1.66−1.59 (m, 3H). LCMS $R_t$ = 0.461 min in 1 min chromatography, purity 88.53%, MS ESI *m/z*: 403.1 $[M+H]^+$.

*7-Amino-5-chloropyrazolo[1,5-a]pyrimidine-3-carbonitrile (**44o**):* Intermediate **44o** was prepared

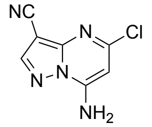


from **43** and ammonium hydroxide in the same manner as described for the preparation of **44a**. Light yellow solid (5.17 g, 2.82 mmol, crude). $^{1}$H NMR (400 MHz, DMSO-$d_6$) $\delta$ 8.64 (s, 1H), 8.48−8.38 (m, 2H), 6.30 (s, 1H). LCMS $R_t$ = 0.415 min in 1 min chromatography, purity 94.13%, MS ESI *m/z*: 194.0 $[M+H]^+$.

*tert-Butyl (5-chloro-3-cyanopyrazolo[1,5-a]pyrimidin-7-yl)carbamate (**49**):* To a solution of

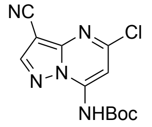


compound **44o** (5.17 g, 26.68 mmol, 1 *eq*) in THF (70 mL) was added DMAP (326 mg, 2.67 mmol, 0.1 *eq*) and $Boc_2O$ (11.65 g, 53.37 mmol, 2 *eq*) at 25°C. Then the mixture was stirred at 50°C. After 3 hours, the reaction mixture was quenched by water (20 mL) and extracted with EtOAc (100 mL×2). The combined organic layers were dried over $Na_2SO_4$, filtered, and concentrated in *vacuo*. The residue was purified by flash silica gel chromatography (eluent: 0-1%, MeOH / DCM) to obtain compound **49** (5.19 g, 16.97 mmol, 96% purity) as a white solid. $^{1}$H NMR (400 MHz, DMSO-$d_6$) $\delta$ 10.98 (s, 1H), 8.83 (s, 1H), 7.59 (s, 1H), 1.53 (s, 9H). LCMS $R_t$ = 0.572 min in 1 min chromatography, purity 97.72%, MS ESI *m/z*: 294.0 $[M+H]^+$.

*tert-Butyl (5-(6-acetamido-1H-indol-1-yl)-3-cyanopyrazolo[1,5-a]pyrimidin-7-yl)carbamate (**50**):*

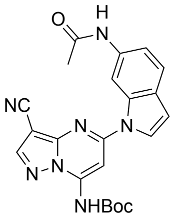


Intermediate **50** was prepared from **49** and **46** in the same manner as described for the preparation of **17**. Light yellow solid (1.50 g, 2.02 mmol, 24.68% yield, 52% purity). LCMS $R_t$ = 1.556 min in 2.5 min chromatography, purity 58.00%, MS ESI *m/z*: 432.3 $[M+H]^+$.

*N-(1-(7-amino-3-cyanopyrazolo[1,5-a]pyrimidin-5-yl)-1H-indol-6-yl)acetamide (**51**):* Compound

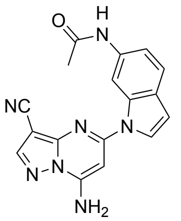


**50** (600 mg, 1.39 mmol, 1 eq) was dissolved in DCM (4 mL) and TFA (0.5 mL). Then the mixture was stirred at 25°C for 2 hours. The reaction mixture was concentrated in *vacuo*. The residue was triturated with MeCN (5 mL). The solid was washed with saturated aqueous $NaHCO_3$ (5 mL×2) and dried in *vacuo*. The crude product was used in the next step directly. Compound **51** (550 mg, 1.15 mmol, 69.15% purity) was obtained as a light brown solid. $^{1}$H NMR (400 MHz, DMSO-$d_6$) $\delta$ 10.02 (s, 1H), 8.59 (s, 2H), 8.37−8.30 (m, 1H), 7.84 (s, 1H), 7.56 (d, $J$ = 8.4 Hz, 1H), 7.38−7.34

(m, 1H), 6.72 (s, 1H), 6.47 (s, 1H), 2.07 (s, 3H). LCMS $R_t$ = 0.463 min in 1 min chromatography, purity 69.15%, MS ESI *m/z*: 332.1 $[M+H]^+$.

*5-Chloro-7-(phenylthio)pyrazolo[1,5-a]pyrimidine-3-carbonitrile (**44q**):* Intermediate **44q** was

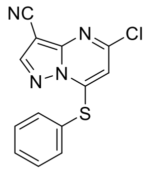

prepared from **43** and ammonium hydroxide in the same manner as described for the preparation of **44a**. White solid (860 mg, 2.82 mmol, 75.18% yield, 94.13% purity). $^1$H NMR (400 MHz, DMSO-$d_6$) $\delta$ 8.94 (m, 1H), 7.81−7.79 (m, 2H), 7.68−7.73 (m, 3H), 6.14 (s, 1H). LCMS $R_t$ = 0.564 min in 1 min chromatography, purity 94.13%, MS ESI *m/z*: 286.9 $[M+H]^+$.

**Synthesis of Compounds (8–33)**

*N-(5-((3-Cyano-7-(cyclopropylamino)pyrazolo[1,5-a]pyrimidin-5-yl)amino)-2-(methyl(2-(methylamino)ethyl)amino)phenyl)propionamide (**8**):* To a solution of compound **42a** (700 mg,

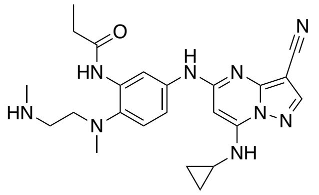

1.28 mmol) in DCM (5 mL) was added TFA (10.74 g, 94.19 mmol) at 25℃. The mixture was stirred at 25°C. After 3 hours, the reaction mixture was quenched by water (10 mL) and extracted with EtOAc (100 mL×2). The combined organic layers were dried over $Na_2SO_4$, filtered, and concentrated in *vacuo*. The residue was purified by prep-HPLC (column: Xtimate C18 150*40mm*10um; mobile phase: [water (FA)-ACN]; B%: 10%-50%, 12min). Then the impure product was purified by prep-HPLC (column: Phenomenex C18 75*30mm*3um; mobile phase: [water ($NH_3H_2O$+$NH_4HCO_3$)-ACN]; B%: 15%-85%, 11min) to obtain **8** (61.9 mg, 11.0%) as a white solid. $^1$H NMR (400 MHz, DMSO-$d_6$) $\delta$ 10.13 (s, 1H), 9.61 (s, 1H), 8.32 (s, 1H), 8.29 (d, *J* = 2.0 Hz, 1H), 8.16 (s, 1H), 7.80 (br d, *J* = 8.0 Hz, 1H), 7.20 (d, *J* = 8.8 Hz, 1H), 6.04 (s, 1H), 2.77−2.74 (m, 2H), 2.64 (s, 3H), 2.58−2.55 (m, 4H), 2.40 (q, *J* = 7.6 Hz, 2H), 2.34 (s, 3H), 1.11 (t, *J* = 7.6 Hz, 3H), 0.80−0.77 (m, 2H), 0.72−0.68 (m, 2H). $^{13}$C NMR (101 MHz, DMSO) $\delta$ 172.0, 157.0, 150.9, 148.2, 145.0, 137.4, 136.2, 134.5, 121.2, 114.9, 114.6, 111.5, 76.2, 58.1, 48.8, 40.8, 39.5, 36.4, 29.6, 23.3, 9.8, 6.5. HPLC Rt = 3.230 min in 8 min chromatography, purity 99.1%. LCMS $R_t$ = 1.201 min in 4 min chromatography, purity 99.5%, MS ESI *m/z:* 448.4 $[M+H]^+$.

*N-(2-((2-Aminoethyl)(methyl)amino)-5-((3-cyano-7-(cyclopropylamino)pyrazolo[1,5-a]pyrimidin-5-yl)amino)phenyl)propionamide (**9**):* Compound **9** was prepared from **42b** in the same manner as described for the preparation of **8**. White solid (25 mg, 17.4%). $^1$H NMR (400 MHz, DMSO-$d_6$) $\delta$ 10.23 (s, 1H), 9.60 (s, 1H), 8.38− 8.27 (m, 2H), 8.23−8.02 (m, 1H), 7.84−7.75 (m, 1H), 7.17 (d, $J$ = 8.8 Hz, 1H), 6.04 (s, 1H), 2.70 (br s, 3H), 2.63 (s, 3H), 2.60−2.55 (m, 2H), 2.45−2.40 (m, 2H), 1.09 (t, $J$ = 7.2 Hz, 3H), 0.81−0.76 (m, 2H), 0.73−0.68 (m, 2H). $^{13}$C NMR (101 MHz, DMSO) $\delta$ 172.1, 157.0, 151.0, 148.2, 145.0, 137.7, 135.9, 134.0, 120.7, 114.9, 114.5, 111.7, 76.2, 60.3, 29.6, 23.3, 9.8, 6.5. HPLC Rt = 2.078 min in 8 min chromatography, purity 99.2%. LCMS $R_t$ = 1.182 min in 4 min chromatography, purity 98.4%, MS ESI *m/z:* 434.4 [M+H]$^+$.

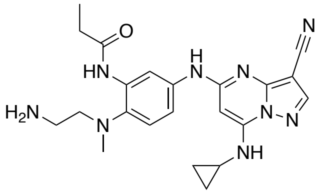

*N-(5-((7-(Cyclopropylamino)-3-ethynylpyrazolo[1,5-a]pyrimidin-5-yl)amino)-2-((2-(ethylamino)ethyl)(methyl)amino)phenyl)propionamide (**10**):* Compound **10** was prepared from **42c** in the same manner as described for the preparation of **8.** Yellow solid (40 mg, 28.0%). $^1$H NMR (400 MHz, DMSO-$d_6$) δ 9.65 (s, 1H), 9.58 (s, 1H), 8.45 (s, 1H), 8.33 (s, 1H), 8.23−8.13 (m, 2H), 7.89−7.79 (m, 1H), 7.19 (d, $J$ = 8.8 Hz, 1H), 6.04 (s, 1H), 3.06−2.97(m, 2H), 2.92−2.78(m, 4H), 2.62−2.57 (m, 1H), 2.54 (s, 3H), 2.53−2.51 (m, 2H), 1.19−1.07 (m, 6H), 0.84−0.76 (m, 2H), 0.75−0.67 (m, 2H). $^{13}$C NMR (101 MHz, DMSO) $\delta$ 172.4, 156.9, 150.9, 148.2, 145.0, 137.7, 136.5, 134.0, 121.2, 115.0, 114.9, 112.6, 76.3, 44.30 41.9, 39.5, 29.6, 23.3, 9.9, 6.5. HPLC Rt = 4.087 min in 8 min chromatography, purity 97.7%. LCMS $R_t$ = 1.955 min in 4 min chromatography, purity 98.7%, MS ESI *m/z:* 462.2 [M+H]$^+$.

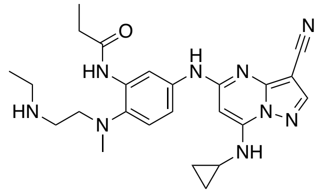

*N-(5-((3-Cyano-7-(cyclopropylamino)pyrazolo[1,5-a]pyrimidin-5-yl)amino)-2-(methyl(2-(pyrrolidin-1-yl)ethyl)amino)phenyl)propionamide (**11**):* To a solution of compound **39d** (408 mg, 1.40 mmol) and compound **41** (255 mg, 1.09 mmol) in dioxane (5 mL) was added $Cs_2CO_3$ (1.07 g, 3.27 mmol), BINAP (102 mg, 0.16 mmol) and $Pd(OAc)_2$ (37 mg, 0.16 mmol) at 25°C. Then the mixture was degassed and purged with $N_2$. The reaction mixture was heated in a microwave at 130°C for 0.5h. The mixture was cooled to room temperature and water (10 mL) was added to the reaction, which was extracted with DCM (4×15 mL). The organic layer was washed with water (2×10 mL), dried with $Na_2SO_4$, filtered, and concentrated to give crude product, which was purified by flash chromatography (eluent of 0~5%, MeOH/DCM) to give the desired product **11** (85 mg, 15.7%) as a white solid. $^1$H NMR (400 MHz, DMSO-$d_6$) $\delta$

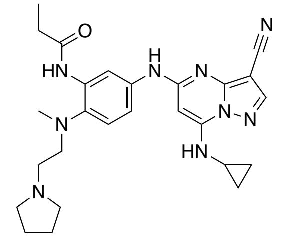

9.65 (s, 1H), 9.57 (s, 1H), 8.33 (s, 1H), 8.19 (br d, *J* = 7.6 Hz, 2H), 7.86 (br d, *J* = 8.0 Hz, 1H), 7.27 (br d, *J* = 8.8 Hz, 1H), 6.05 (s, 1H), 2.88 (br s, 2H), 2.64 (s, 3H), 2.58 (br s, 2H), 2.47 (br s, 2H), 2.44−2.29 (m, 5H), 1.75−1.66 (m, 4H), 1.11 (t, *J* = 7.6 Hz, 3H), 0.83−0.76 (m, 2H), 0.73−0.67 (m, 2H). $^{13}$C NMR (101 MHz, DMSO) $\delta$ 171.5, 157.0, 150.9, 148.2, 145.0, 122.2, 122.2, 114.9, 111.4, 76.2, 56.7, 53.9, 43.0, 39.5, 29.7, 23.1, 9.8, 6.5. HPLC Rt = 3.712 min in 8 min chromatography, purity 99.1%. LCMS $R_t$ = 1.280 min in 4 min chromatography, purity 99.0%, MS ESI *m/z:* 488.5 $[M+H]^+$.

*N-(5-((3-Cyano-7-(cyclopropylamino)pyrazolo[1,5-a]pyrimidin-5-yl)amino)-2-(methyl(2-(piperidin-1-yl)ethyl)amino)phenyl)propionamide (**12**):* Compound **12** was prepared from **39e** and **41** in the same manner as described for the preparation of compound **11.** Yellow solid (37.7 mg, 14.0%). $^{1}$H NMR (400 MHz, DMSO-$d_6$) $\delta$ 9.65 (s, 1H), 9.17 (s, 1H), 8.34 (s, 1H), 8.19−8.13 (m, 2H), 7.85 (d, *J* = 7.6 Hz, 1H), 7.25 (d, *J* = 8.8 Hz, 1H), 6.05 (s, 1H), 2.92 (t, *J* = 6.0 Hz, 2H), 2.62 (s, 3H), 2.46−2.40 (m, 3H), 2.35−2.27 (m, 6H), 1.50−1.46 (m, 4H), 1.41−1.35 (m, 2H), 1.13 (t, *J* = 7.6 Hz, 3H), 0.81−0.79 (m, 2H), 0.73−0.71 (m, 2H).

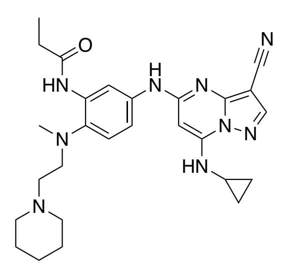

$^{13}$C NMR (101 MHz, DMSO) $\delta$ 171.5, 156.9, 150.9, 148.2, 145.0, 137.2, 136.6, 134.4, 121.9, 115.1, 114.8, 111.9, 76.3, 76.3, 56.5, 54.3, 43.0, 30.0, 25.3, 24.0, 23.3, 9.8, 6.5. HPLC Rt = 3.510 min in 8 min chromatography, purity 99.2%. LCMS $R_t$ = 1.319 min in 4 min chromatography, purity 99.5%, MS ESI *m/z:* 502.5 $[M+H]^+$.

*N-(2-(3-Aminopiperidin-1-yl)-5-((3-cyano-7-(cyclopropylamino)pyrazolo[1,5-a]pyrimidin-5-yl)amino)phenyl)propionamide (**13**):* Compound **13** was prepared from **42f** in the same manner as described for the preparation of **8.** White solid (67.6 mg, 13.5%). $^{1}$H NMR (400 MHz, DMSO-$d_6$) $\delta$ 9.57 (s, 1H), 8.94 (s, 1H), 8.29 (s, 1H), 8.09 (br d, *J* = 1.2 Hz, 1H), 7.74 (br d, *J* = 7.6 Hz, 1H), 7.05 (d, *J* = 8.8 Hz, 1H), 5.99 (s, 1H), 3.39−3.28 (m, 2H), 2.89−2.85 (m, 1H), 2.76−2.74 (m, 2H), 2.65-2.59 (m, 1H), 2.55−2.54 (m, 1H), 2.47−2.37 (m, 3H), 1.79−1.78 (m, 3H), 1.775−1.67 (m, 1H), 1.25−1.17 (m, 1H), 1.09 (t, *J* = 7.6 Hz, 3H), 0.76−0.73 (m, 2H), 0.68−0.65 (m, 2H). $^{13}$C NMR (101 MHz, DMSO) $\delta$ 171.7, 157.0, 150.9, 148.2, 145.0, 137.9, 136.0, 132.9, 120.2, 115.0, 114.9, 112.2, 76.3, 61.4, 52.1, 47.6, 32.8, 29.9, 23.3, 9.8, 6.6. HPLC Rt = 3.443 min in 8 min chromatography, purity 98.4%. LCMS $R_t$ = 1.286 min in 4 min chromatography, purity 98.4%, MS ESI *m/z:* 460.4 $[M+H]^+$.

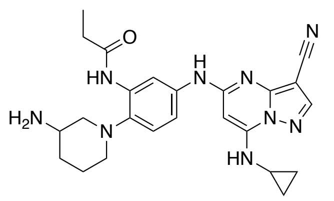

*N-(5-((3-Cyano-7-(cyclopropylamino)pyrazolo[1,5-a]pyrimidin-5-yl)amino)-2-(piperazin-1-yl)phenyl)propionamide (**14**):* Compound **14** was prepared from **42g** and **41** in the same manner

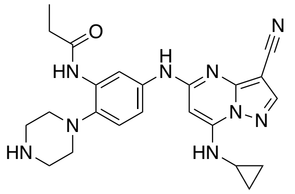

as described for the preparation of **8.** White solid (106.8 mg, 28.7%). $^1$H NMR (400 MHz, DMSO-$d_6$) $\delta$ 9.64 (s, 1H), 8.82 (s, 1H), 8.33 (s, 1H), 8.18 (s, 1H), 8.09 (s, 1H), 7.82 (br d, $J$ = 8.0 Hz, 1H), 7.14 (d, $J$ = 8.8 Hz, 1H), 6.03 (s, 1H), 3.34−3.14 (m, 1H), 2.89−2.87 (m, 4H), 2.72−2.69 (m, 4H), 2.60−2.56 (m, 1H), 2.42 (q, $J$ = 7.6 Hz, 2H), 1.13 (t, $J$ = 7.2 Hz, 3H), 0.80−0.78 (m, 2H), 0.73−0.69 (m, 2H). $^{13}$C NMR (101 MHz, DMSO) $\delta$ 171.5, 156.9, 150.9, 148.2, 145.0, 137.9, 136.3, 132.9, 120.3, 115.1, 114.9, 112.3, 76.3, 76.2, 53.1, 46.0, 30.1, 23.3, 9.8, 6.5. HPLC Rt = 2.952 min in 8 min chromatography, purity 98.2%. LCMS $R_t$ = 1.168 min in 4 min chromatography, purity 98.0%, MS ESI *m/z:* 446.4 [M+H]$^+$.

*N-(5-((3-Cyano-7-(cyclopropylamino)pyrazolo[1,5-a]pyrimidin-5-yl)amino)-2-(3-(methylamino)piperidin-1-yl)phenyl)propionamide (**15**):* Compound **15** was prepared from **42h**

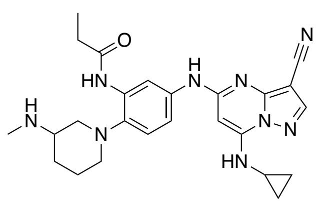

and **41** in the same manner as described for the preparation of **8.** White solid (59.3 mg, 14.2%). $^1$H NMR (400 MHz, DMSO-$d_6$) $\delta$ 9.61 (s, 1H), 8.99 (s, 1H), 8.32 (s, 1H), 8.16−8.13 (m, 2H), 7.78 (d, $J$ = 6.8 Hz, 1H), 7.09 (d, $J$ = 8.8 Hz, 1H), 6.03 (s, 1H), 2.882.80 (m, 2H), 2.68−2.58 (m, 4H), 2.45−2.40 (m, 3H), 2.30 (s, 3H), 1.76−1.74 (m, 2H), 1.65−1.59 (m, 1H), 1.36−1.29 (m, 1H), 1.13 (t, $J$ = 7.6 Hz, 3H), 0.81−0.77 (m, 2H), 0.75−0.63 (m, 2H). $^{13}$C NMR (101 MHz, DMSO) $\delta$ 171.8, 157.0, 151.0, 148.2, 145.0, 138.0, 136.1, 133.0, 120.3, 115.1, 114.9, 112.3, 76.3, 76.21, 58.1, 55.7, 52.3, 33.8, 29.9, 29.1, 23.3, 23.3, 9.9, 6.6. HPLC Rt = 3.201 min in 8 min chromatography, purity 97.5%. LCMS $R_t$ = 1.306 min in 4 min chromatography, purity 98.3%, MS ESI *m/z:* 474.4 [M+H]$^+$.

*N-(1-(3-cyano-7-(cyclopropylamino)pyrazolo[1,5-a]pyrimidin-5-yl)-1H-indol-5-yl)acetamide (16):*

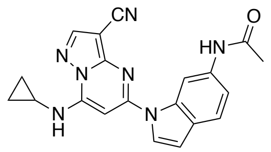

The compound **16** was previously characterized as compound **29** in Reference [2].

*N-(1-(3-Cyano-7-(cyclobutylamino)pyrazolo[1,5-a]pyrimidin-5-yl)-1H-indol-6-yl)acetamide (**17**):*

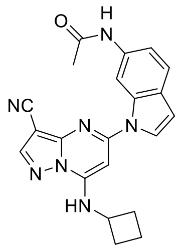

To a solution of **44a** (200 mg, 0.81 mmol) and **46** (169 mg, 0.96 mmol) in dioxane (5 mL) was added $Cs_2CO_3$ (789 mg, 2.42 mmol), Xantphos (70 mg, 0.12 mmol) and $Pd(OAc)_2$ (27 mg, 0.12 mmol) at 25°C. The mixture was purged with $N_2$ and heated in a microwave condition at 130°C for 0.5 h. The mixture was cooled to rt and water (10 mL) was added to the reaction, which was extracted with DCM (3×15 mL). The organic layer was washed with water (2×10 mL), dried with $Na_2SO_4$, filtered, and concentrated to give crude product, which was purified by prep-HPLC (column: Phenomenex Gemini-NX C18 75*30mm*3um; mobile phase: [water (0.05%$NH_3H_2O$+10mM $NH_4HCO_3$)-ACN]; B%: 36%-66%, 10min) to get **17** (74.1 mg, 0.19 mmol, 23.62% yield, 99.22% purity) as a yellow solid. $^{1}$H NMR (DMSO-$d_6$, 400 MHz) $\delta$ 10.00 (s, 1H), 8.84 (s, 1H), 8.75 (s, 1H), 8.63 (s, 1H), 8.05 (d, $J$ = 3.6 Hz, 1H), 7.54 (d, $J$ = 8.4 Hz, 1H), 7.28 (dd, $J$ = 1.6, 8.4 Hz, 1H), 6.72 (d, $J$ = 3.6 Hz, 1H), 6.59−6.46 (m, 1H), 4.45 4.40 (m, 1H), 2.47−2.37 (m, 2H), 2.35−2.22 (m, 2H), 2.08 (s, 3H), 1.82−1.66 (m, 2H). $^{13}$C NMR (101 MHz, DMSO) $\delta$ 168.0, 153.7, 150.0, 147.4, 146.5, 135.5, 134.8, 126.7, 126.4, 120.7, 115.4, 114.0, 106.6, 106.0, 79.0, 79.0, 46.8, 29.3, 24.0, 14.8. HPLC Rt = 3.101 min in 8 min chromatography, XBridge Shield RP18 (2.1×50mm 5μm), purity 99.22%. LCMS $R_t$ = 1.604 min in 4 min chromatography, Xtimate C18,3um,2.1*30mm, purity 97.76%, MS ESI *m/z:* 386.4 [M+H]$^+$.

*N-(1-(3-Cyano-7-((3,3-difluorocyclobutyl)amino)pyrazolo[1,5-a]pyrimidin-5-yl)-1H-indol-6-yl)acetamide (**18**):* Compound **18** was prepared from **44b** and **46** in the same manner as

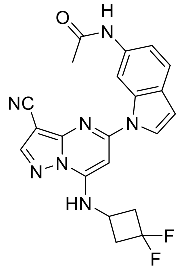

described for the preparation of **17**. White solid (69.7 mg, 0.16 mol, 31.12% yield, 99.49% purity). $^{1}$H NMR (400 MHz, DMSO-$d_6$) $\delta$ 9.99 (s, 1H), 9.04 (br s, 1H), 8.84 (s, 1H), 8.62 (s, 1H), 8.08 (d, $J$ = 3.6 Hz, 1H), 7.53 (d, $J$ = 8.0 Hz, 1H), 7.28 (dd, $J$ = 1.2, 8.4 Hz, 1H), 6.74 (d, $J$ = 3.6 Hz, 1H), 6.56 (s, 1H), 4.44−4.30 (m, 1H), 3.27−3.11 (m, 2H), 2.99 (m, 2H), 2.06 (s, 3H). $^{13}$C NMR (101 MHz, DMSO) $\delta$ 168.0, 153.8, 150.0, 148.1, 146.4, 135.4, 134.9, 126.8, 126.4, 120.6, 115.5, 114.1, 106.5, 106.3, 79.3, 78.6, 42.06, 41.8, 41.6, 24.0. HPLC Rt = 3.179 min in 8 min chromatography, Xtimate C18 2.1*30mm 3um, purity 99.67%. LCMS $R_t$ = 1.943 min in 4 min chromatography, Xtimate C18,3um,2.1*30mm, purity 99.49%, MS ESI *m/z*: 422.0 [M+H]$^+$.

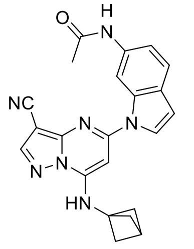

*N-(1-(7-(Bicyclo[1.1.1]pentan-1-ylamino)-3-cyanopyrazolo[1,5-a]pyrimidin-5-yl)-1H-indol-6-yl)acetamide (**19**):* Compound **19** was prepared from **44c** and **46** in the same manner as described for the preparation of **17**. White solid (41.4 mg, 0.10 mmol, 13.33% yield, 98.51% purity). **$^1$H NMR** (400 MHz, DMSO-$d_6$) $\delta$ 10.01 (s, 1H), 9.29 (br s, 1H), 8.79 (s, 1H), 8.62 (s, 1H), 7.95 (d, $J$ = 3.6 Hz, 1H), 7.56 (d, $J$ = 8.4 Hz, 1H), 7.22 (dd, $J$ = 1.6, 8.4 Hz, 1H), 6.74 (d, $J$ = 3.6 Hz, 1H), 6.56 (s, 1H), 2.55 (s, 1H), 2.33−2.23 (m, 6H), 2.07 (s, 3H). $^{13}$C NMR (101 MHz, DMSO) $\delta$ 168.0, 153.1, 150.0, 147.6, 146.5, 135.6, 134.7, 126.7, 126.4, 120.9, 115.2, 114.1, 106.5, 104.7, 80.0, 78.8, 52.5, 49.6, 24.2, 24.0. HPLC Rt = 3.316 min in 8 min chromatography, Xtimate C18 2.1*30mm 3um, purity 98.514%. LCMS $R_t$ = 1.642 min in 4 min chromatography, Xtimate C18, 3um, 2.1*30mm, purity 98.64%, MS ESI *m/z*: 398.3 [M+H]$^+$.

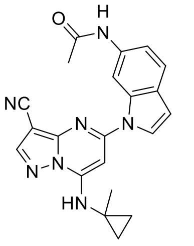

*N-(1-(3-Cyano-7-((1-methylcyclopropyl)amino)pyrazolo[1,5-a]pyrimidin-5-yl)-1H-indol-6-yl)acetamide (**20**):* Compound **20** was prepared from **44d** and **46** in the same manner as described for the preparation of **17**. White solid (14 mg, 0.4 mmol, 3.49% yield, 99.66 % purity). $^1$H NMR (DMSO-$d_6$, 400 MHz) $\delta$ 10.01 (s, 1H), 9.03 (br s, 1H), 8.84 (s, 1H), 8.62 (s, 1H), 8.02 (d, $J$ = 3.6 Hz, 1H), 7.58 (d, $J$ = 0.4 Hz, 1H), 7.20 (dd, $J$ = 1.2, 8.4 Hz, 1H), 6.76 (d, $J$ = 3.6 Hz, 1H), 6.68 (s, 1H), 2.08 (s, 3H), 1.49 (s, 3H), 1.04−0.97 (m, 2H), 0.96−0.88 (m, 2H). $^{13}$C NMR (101 MHz, DMSO) $\delta$ 167.9, 153.1, 150.0, 148.7, 146.6, 135.6, 134.6, 126.6, 126.5, 120.9, 115.1, 114.0, 106.4, 104.8, 80.2, 78.7, 29.8, 24.0, 21.2, 14.2. HPLC Rt = 3.094 min in 8 min chromatography, Xtimate C18 2.1*30mm 3um, purity 99.467%. LCMS $R_t$ = 1.847 min in 4 min chromatography, Xtimate C18,3um,2.1*30mm, purity 99.792%, MS ESI *m/z*: 386.1 [M+H]$^+$.

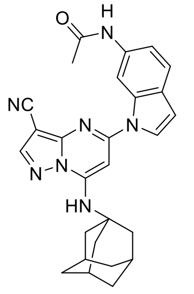

*N-(1-(7-(((3s,5s,7s)-Adamantan-1-yl)amino)-3-cyanopyrazolo[1,5-a]pyrimidin-5-yl)-1H-indol-6-yl)acetamide (**21**):* Compound **21** was prepared from **44e** and **46** in the same manner as described for the preparation of **17**. White solid (24.8 mg, 0.05 mmol, 6.69% yield, 99.56 % purity). $^1$H NMR (DMSO-$d_6$, 400 MHz) $\delta$ 10.02 (s, 1H), 8.85 (s, 1H), 8.65 (s, 1H), 8.00 (d, $J$ = 3.6 Hz, 1H), 7.57 (d, $J$ = 8.0 Hz, 1H), 7.17 (dd, $J$ = 1.6, 8.4 Hz, 1H), 7.10 (s, 1H), 6.75 (d, $J$ = 3.2 Hz, 1H), 6.66 (s, 1H), 2.16 (s, 9H), 2.05 (s, 3H), 1.79−1.60 (m, 6H). $^{13}$C NMR (101 MHz, DMSO) $\delta$ 167.9, 152.7, 149.6, 146.4, 146.1, 135.6, 134.6, 126.9, 126.4, 121.0, 115.1, 113.8, 106.5, 104.5, 81.5, 79.3, 53.2, 40.7, 35.3, 28.9, 23.9. HPL**C** Rt = 4.399 min in 8 min chromatography, Xtimate C18 2.1*30mm 3um, purity 99.56%. LCMS $R_t$ = 2.611 min in 4 min chromatography, Xtimate C18, 3um, 2.1*30mm, purity 99.55%, MS ESI *m/z*: 466.2 [M+H]$^+$.

*N-(1-(7-(((1r,3r,5r,7r)-Adamantan-2-yl)amino)-3-cyanopyrazolo[1,5-a]pyrimidin-5-yl)-1H-indol-6-yl)acetamide (**22**):* Compound **22** was prepared from **44f** and **46** in the same manner as

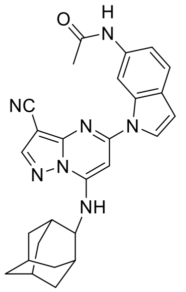

described for the preparation of **17**. White solid (38.4 mg, 0.08 mmol, 13.42% yield, 99.29 % purity). $^1$H NMR (DMSO-$d_6$, 400 MHz) *δ* 10.01 (s, 1H), 8.87 (s, 1H), 8.66 (s, 1H), 8.05 (s, 1H), 7.59−7.42 (m, 1H), 7.32−7.01 (m, 2H), 6.81−6.49 (m, 2H), 4.21−7.17 (m, 1H), 2.16 (s, 2H), 2.07 (s, 3H), 1.99−1.95 (m, 4H), 1.89−1.84 (m, 4H), 1.74 (s, 2H), 1.65−1.61 (m, 2H). $^{13}$C NMR (101 MHz, DMSO) *δ* 167.9, 153.77, 149.6, 147.2, 146.8, 135.5, 134.8, 126.7, 126.4, 120.7, 115.4, 113.9, 106.6, 105.9, 79.5, 79.1, 55.26, 36.8, 36.1, 31.1, 30.8, 26.5, 24.0. HPLC Rt = 4.328 min in 8 min chromatography, XBridge Shield RP18 (2.1×50mm 5μm), purity 99.29%. LCMS $R_t$ = 2.238 min in 4 min chromatography, Xtimate C18,3um,2.1*30mm, purity 99.88%, MS ESI *m/z*: 466.4 $[M+H]^+$.

*N-(1-(3-Cyano-7-(phenylamino)pyrazolo[1,5-a]pyrimidin-5-yl)-1H-indol-6-yl)acetamide (**23**):*

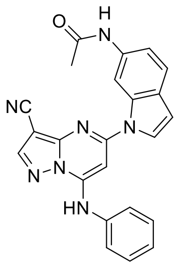

Compound **23** was prepared from **44g** and **46** in the same manner as described for the preparation of **17**. White solid (10.4 mg, 0.02 mmol, 8.67% yield, 99.75% purity). $^1$H NMR (400 MHz, DMSO-$d_6$) *δ* 10.94−10.25 (m, 1H), 10.02 (s, 1H), 8.76 (s, 1H), 8.74 (s, 1H), 7.81 (d, *J* = 3.6 Hz, 1H), 7.65−7.46 (m, 5H), 7.38−7.28 (m, 1H), 7.20 (dd, *J* = 1.2, 8.4 Hz, 1H), 6.69 (d, *J* = 3.2 Hz, 1H), 6.58 (s, 1H), 2.11 (s, 3H). $^{13}$C NMR (101 MHz, DMSO) δ 168.6, 154.0, 150.8, 147.8, 147.4, 136.9, 136.1, 135.1, 130.2, 127.0, 126.8, 126.7, 124.9, 121.4, 115.6, 114.4, 107.2, 105.0, 81.4, 79.7, 24.5. HPLC Rt = 3.200 min in 8 min chromatography, Xtimate C18 2.1*30mm 3um, purity 99.75%. LCMS $R_t$ = 1.939 min in 4 min chromatography, Xtimate C18,3um,2.1*30mm, purity 97.91%, MS ESI *m/z*: 408.0 $[M+H]^+$.

*N-(1-(3-Cyano-7-((oxetan-3-ylmethyl)amino)pyrazolo[1,5-a]pyrimidin-5-yl)-1H-indol-6-yl)acetamide (**24**):* Compound **24** was prepared from **44h** and **46** in the same manner as

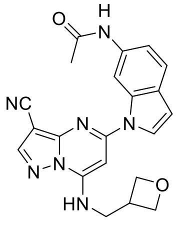

described for the preparation of **17**. White solid (27.7 mg, 0.68 mmol, 9.06% yield, 99.56% purity). $^1$H NMR (400 MHz, DMSO-$d_6$) *δ* 10.00 (s, 1H), 8.86 (s, 1H), 8.82 (s, 1H), 8.61 (s, 1H), 8.05 (d, *J* = 3.6 Hz, 1H), 7.53 (d, *J* = 8.4 Hz, 1H), 7.28 (d, *J* = 8.8 Hz, 1H), 6.73 (d, *J* = 3.6 Hz, 1H), 6.66 (s, 1H), 4.69 (m, 2H), 4.41 (t, *J* = 6.0 Hz, 2H), 3.84 (d, *J* = 7.6 Hz, 2H), 3.48−3.39 (m, 1H), 2.08 (s, 3H). $^{13}$C NMR (101 MHz, DMSO) *δ* 168.1, 153.9, 149.9, 148.6, 146.6, 135.4, 134.8, 126.8, 126.4, 120.6, 115.5, 114.1, 106.6, 106.3, 78.8, 78.6, 74.0, 44.3, 33.4, 24.0, 1.2. HPLC Rt = 2.474 min in 8 min chromatography, Xtimate C18 2.1*30mm 3um, purity 99.563%. LCMS $R_t$ = 1.615 min in 4 min chromatography, Xtimate C18,3um,2.1*30mm, purity 99.505%, MS ESI *m/z*: 402.0 $[M+H]^+$.

*N-(1-(3-Cyano-7-(((tetrahydrofuran-3-yl)methyl)amino)pyrazolo[1,5-a]pyrimidin-5-yl)-1H-indol-6-yl)acetamide (**25**):* Compound **25** was prepared from **44i** and **46** in the same manner as

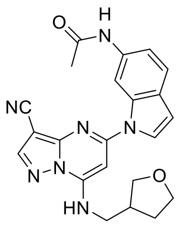

described for the preparation of **17**. White solid (32.2 mg, 0.08 mmol, 10.65% yield, 98.94 % purity). $^1$H NMR (DMSO-$d_6$, 400 MHz) $\delta$ 10.00 (s, 1H), 8.86 (s, 1H), 8.79 (s, 1H), 8.64 (s, 1H), 8.07 (d, $J$ = 3.6 Hz, 1H), 7.55 (d, $J$ = 8.4 Hz, 1H), 7.27 (dd, $J$ = 1.6, 8.4 Hz, 1H), 6.75 (d, $J$ = 3.6 Hz, 1H), 6.68 (s, 1H), 3.83−3.79 (m, 1H), 3.75−3.71 (m, 1H), 3.65−3.61 (m, 1H), 3.54 (s, 3H), 2.82−2.62 (m, 1H), 2.07 (s, 3H), 2.05−1.93 (m, 1H), 1.76−1.63 (m, 1H). $^{13}$C NMR (101 MHz, DMSO) $\delta$ 168.0, 153.8, 149.9, 148.6, 146.6, 135.4, 134.8, 126.8, 126.4, 120.7, 115.4, 114.0, 106.5, 106.1, 78.7, 78.7, 70.4, 66.8, 44.3, 37.8, 29.4, 24.0. HPLC Rt = 2.690 min in 8 min chromatography, Xtimate C18 2.1*30mm 3um, purity 98.94%. LCMS $R_t$ = 1.356 min in 4 min chromatography, Xtimate C18,3um,2.1*30mm, purity 98.99%, MS ESI *m/z*: 416.4 $[M+H]^+$.

*N-(1-(3-Cyano-7-(((tetrahydro-2H-pyran-4-yl)methyl)amino)pyrazolo[1,5-a]pyrimidin-5-yl)-1H-indol-6-yl)acetamide (**26**):* Compound **26** was prepared from **44j** and **46** in the same manner as

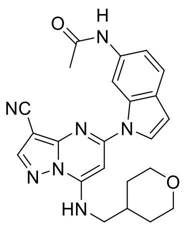

described for the preparation of **17**. White solid (47.9 mg, 0.1 mmol, 13.64% yield, 96.45 % purity). $^1$H NMR (DMSO-$d_6$, 400 MHz) $\delta$ 10.01 (s, 1H), 8.84 (s, 1H), 8.61 (s, 1H), 8.05 (d, $J$ = 3.2 Hz, 1H), 7.55 (d, $J$ = 8.4 Hz, 1H), 7.25 (dd, $J$ = 1.2, 8.4 Hz, 1H), 6.74 (d, $J$ = 3.2 Hz, 1H), 6.62 (s, 1H), 3.84 (dd, $J$ = 2.4, 10.8 Hz, 2H), 3.45−3.39 (m, 2H), 3.32−3.20 (m, 3H), 2.07 (s, 3H), 2.06−1.96 (m, 1H), 1.68−1.61 (m, 2H), 1.36−1.25 (m, 2H). $^{13}$C NMR (101 MHz, DMSO) $\delta$ 168.0, 153.7, 150.0, 148.7, 146.5, 135.4, 134.8, 126.8, 126.4, 120.7, 115.3, 114.2, 106.4, 106.0, 78.8, 78.6, 66.6, 47.3, 34.3, 30.2, 24.0. HPLC Rt = 2.785 min in 8 min chromatography, Xtimate C18 2.1*30mm 3um, purity 96.45%. LCMS $R_t$ = 1.418 min in 4 min chromatography, Xtimate C18,3um,2.1*30mm, purity 96.37%, MS ESI *m/z*: 430.4 $[M+H]^+$.

*N-(1-(3-Cyano-7-((1-methylpiperidin-4-yl)amino)pyrazolo[1,5-a]pyrimidin-5-yl)-1H-indol-6-yl)acetamide (**27**):* Compound **27** was prepared from **44k** and **46** in the same manner as

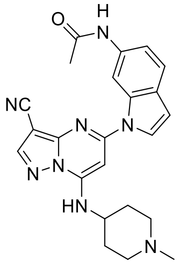

described for the preparation of **17**. White solid (114.9 mg, 0.27 mmol, 38.57% yield, 99.40% purity). $^1$H NMR (400 MHz, DMSO-$d_6$) $\delta$ 10.01 (s, 1H), 8.88 (s, 1H), 8.61 (s, 1H), 8.35 (br s, 1H), 8.07 (d, $J$ = 3.2 Hz, 1H), 7.53 (d, $J$ = 8.4 Hz, 1H), 7.25 (dd, $J$ = 1.6, 8.0 Hz, 1H), 6.72 (d, $J$ = 3.2 Hz, 1H), 6.65 (s, 1H), 3.90−3.72 (m, 1H), 2.79−2.74 (m, 2H), 2.17 (s, 3H), 2.13−2.03 (m, 5H),

1.93−1.76 (m, 4H). $^{13}C$ NMR (101 MHz, DMSO) *δ* 168.0, 153.8, 150.1, 147.7, 146.4, 135.4, 134.8, 126.8, 126.4, 120.6, 115.3, 114.1, 106.3, 106.0, 78.7, 78.6, 54.3, 49.3, 46.0, 30.8, 24.0. HPLC Rt = 2.931 min in 8 min chromatography, Xtimate C18 2.1*30mm 3um, purity 99.40%. LCMS $R_t$ = 1.406 min in 4 min chromatography, Xtimate C18,3um,2.1*30mm, purity 99.49%, MS ESI *m/z*: 429.4 $[M+H]^+$.

*N-(1-(7-((1-Acetylpiperidin-4-yl)amino)-3-cyanopyrazolo[1,5-a]pyrimidin-5-yl)-1H-indol-6-yl)acetamide (**28**):* Compound **28** was prepared from **44l** and **46** in the same manner as

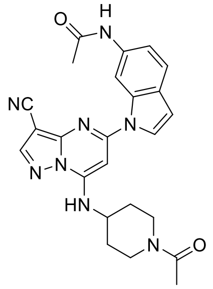

described for the preparation of **17**. White solid (114.9 mg, 0.27 mmol, 38.57% yield, 99.40% purity). $^1H$ NMR (400 MHz, DMSO-$d_6$) *δ* 10.02 (s, 1H), 8.90 (s, 1H), 8.65 (s, 1H), 8.51−8.23 (m, 1H), 8.07 (d, *J* = 3.6 Hz, 1H), 7.56 (d, *J* = 8.4 Hz, 1H), 7.24 (d, *J* = 8.4 Hz, 1H), 6.84−6.68 (m, 2H), 4.47 (br d, *J* = 12.4 Hz, 1H), 4.20−4.07 (m, 1H), 3.90 (br d, *J* = 12.4 Hz, 1H), 3.28−3.15 (m, 1H), 2.76−2.64 (m, 1H), 2.07 (s, 3H), 2.03 (s, 3H), 2.01−1.55 (m, 4H). $^{13}C$ NMR (101 MHz, DMSO) *δ* 168.1, 153.9, 150.1, 147.7, 146.6, 135.5, 134.8, 126.7, 126.4, 120.7, 115.3, 114.0, 106.5, 106.0, 78.95, 78.7, 49.4, 44.8, 31.3, 30.6, 24.3, 21.3. HPLC Rt = 2.583 min in 8 min chromatography, Xtimate C18 2.1*30mm 3um, purity 99.23%. LCMS $R_t$ = 1.689 min in 4 min chromatography, Xtimate C18, 3um, 2.1*30mm, purity 98.42%, MS ESI *m/z*: 457.0 $[M+H]^+$.

*N-(1-(3-Cyano-7-(((1-methyl-1H-pyrazol-4-yl)methyl)amino)pyrazolo[1,5-a]pyrimidin-5-yl)-1H-indol-6-yl)acetamide (**29**):* Compound **29** was prepared from **44m** and **46** in the same manner

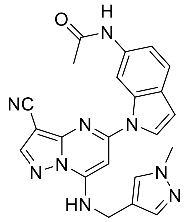

as described for the preparation of **17**. White solid (21.8 mg, 0.05 mmol, 7.04% yield, 95.53 % purity). $^1H$ NMR (DMSO-$d_6$, 400 MHz) *δ* 10.00 (s, 1H), 9.02 (s, 1H), 8.78 (s, 1H), 8.62 (s, 1H), 8.03 (d, *J* = 3.6 Hz, 1H), 7.72 (s, 1H), 7.56 (d, *J* = 8.4 Hz, 1H), 7.50 (s, 1H), 7.32 (dd, *J* = 1.2, 8.4 Hz, 1H), 6.76 (d, *J* = 3.6 Hz, 1H), 6.64 (s, 1H), 4.61 (s, 2H), 3.76 (s, 3H), 2.09 (s, 3H). $^{13}C$ NMR (101 MHz, DMSO) *δ* 168.1, 153.6, 150.0, 148.3, 146.5, 138.2, 135.4, 134.8, 129.6, 126.7, 126.4, 120.7, 117.3, 115.5, 114.1, 106.6, 106.0, 78.7, 78.6, 38.5, 36.2, 24.0. HPLC Rt = 2.496 min in 8 min chromatography, Xtimate C18 2.1*30mm 3um, purity 95.52%. LCMS $R_t$ = 1.277 min in 4 min chromatography, Xtimate C18,3um, 2.1*30mm, purity 96.66%, MS ESI *m/z*: 426.4 $[M+H]^+$.

*(R)-N-(1-(3-Cyano-7-((1-cyanoethyl)amino)pyrazolo[1,5-a]pyrimidin-5-yl)-1H-indol-6-yl)acetamide (**30**):* To a solution of compound **48** (150 mg, 0.37 mmol, 1 eq) in DCM (3 mL) was added TEA (75 mg, 0.75 mmol, 2 eq) at 25°C. And then trifluoroacetic anhydride (TFAA) (156 mg, 0.75 mmol, 2 *eq*) was added to the reaction mixture at 0°C. The mixture was stirred at 25°C for 2 hours. After the starting material was consumed, the reaction mixture was concentrated, and the residue was dissolved in MeOH (5 mL). Then $K_2CO_3$ (200 mg) was added to the mixture. The suspension was stirred at 25℃ for 10 hours. The reaction mixture was filtered. The filter cake was washed with MeOH (3×5 mL) and the solid was dried in vacuo. The residue was purified by prep-HPLC (column: Phenomenex luna 30*30mm*10um+YMC AQ 100*30*10um; mobile phase: [water (0.05% $NH_3H_2O$)-ACN]; B%:30%-60%, 20min) to get compound **30** (17.1 mg, 0.04 mmol, 11.88% yield, 99.57% purity) as a white solid. $^1$H NMR (400 MHz, DMSO-$d_6$) $\delta$ 9.99 (s, 1H), 9.24 (br s, 1H), 8.90 (s, 1H), 8.71 (s, 1H), 8.08 (d, $J$ = 3.6 Hz, 1H), 7.55 (d, $J$ = 8.4 Hz, 1H), 7.36 (dd, $J$ = 1.2, 8.4 Hz, 1H), 6.91 (s, 1H), 6.81 (d, $J$ = 3.2 Hz, 1H), 5.45−5.28 (m, 1H), 2.07 (s, 3H), 1.74 (d, $J$ = 7.2 Hz, 3H). $^{13}$C NMR (101 MHz, DMSO) $\delta$ 167.9, 154.3, 149.8, 147.4, 147.0, 135.6, 135.0, 126.6, 126.5, 120.6, 119.4, 116.1, 113.7, 107.3, 107.2, 79.7, 79.3, 24.0, 18.3. HPLC Rt = 3.791 min in 8 min chromatography, purity 99.57%. LCMS $R_t$ = 1.605 min in 4 min chromatography, purity 99.41%, MS ESI *m/z*: 385.3 [M+H]$^+$.

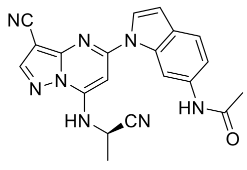


*N-(1-(3-Cyano-7-(phenylsulfonamido)pyrazolo[1,5-a]pyrimidin-5-yl)-1H-indol-6-yl)acetamide (**31**):* To a solution of intermediate **51** (210 mg, 0.57 mmol, 1 eq, HCl) in DCM (3 mL) and pyridine (5 mL) was added TEA (288 mg, 2.85 mmol, 5 eq) and benzenesulfonyl chloride ((302 mg, 1.71 mmol, 3 eq) dropwise at 0℃. Then the mixture was stirred at 25°C for 2 hours. The reaction mixture was concentrated in vacuo. The residue was purified by prep-HPLC (column: Phenomenex Gemini-NX C18 75*30mm*3um; mobile phase: [water (10mM $NH_4HCO_3$)-ACN]; B%: 30%-60%, 20min). The final compound **31** (18.3 mg, 0.037 mmol, 6.58% yield, 96.84% purity) was obtained as a brown solid. $^1$H NMR (400 MHz, DMSO-$d_6$) $\delta$ 9.97 (s, 1H), 8.50 (s, 1H), 8.43 (s, 1H), 7.99−7.91 (m, 2H), 7.59 (d, $J$ = 3.6 Hz, 1H), 7.56−7.48 (m, 4H), 7.35 (dd, $J$ = 1.2, 8.4 Hz, 1H), 6.85 (s, 1H), 6.73 (d, $J$ = 3.6 Hz, 1H), 2.06 (s, 3H). $^{13}$C NMR (101 MHz, DMSO) $\delta$ 167.9, 153.1, 150.9, 150.4, 145.8, 143.0, 135.3, 134.8, 131.8, 128.9, 126.3, 126.0, 120.7, 115.9, 114.4, 106.6, 106.0, 85.7, 78.2, 23.9. HPLC Rt = 4.016 min in 8 min chromatography, purity 95.82%. LCMS $R_t$ = 1.685 min in 4 min chromatography, purity 96.84%, MS ESI *m/z*: 472.1 [M+H]$^+$.

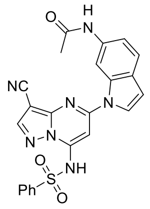

*N-(1-(3-Cyano-7-(methylsulfonamido)pyrazolo[1,5-a]pyrimidin-5-yl)-1H-indol-6-yl)acetamide (**32**):* To a solution of intermediate **51** (100 mg, 0.30 mmol, 1 eq) in DCM (5 mL) was added

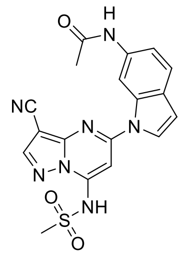

DMAP (4 mg, 0.03 mmol, 0.1 eq), TEA (458 mg, 4.53 mmol, 15 eq) and MsCl (2.14 g, 18.68 mmol, 61.9 eq) at 0℃. Then the mixture was stirred at 25°C for 10 hours. The mixture was quenched with saturated $NaHCO_3$ (10 mL), which was extracted with DCM (2× 10 mL). The organic layer was washed with water (10 mL), dried with $Na_2SO_4$, filtered, and concentrated to give crude product, which was purified by prep-HPLC (column: Phenomenex Gemini-NX C18 75*30mm*3um; mobile phase: [water (10mM $NH_4HCO_3$)-ACN]; B%: 5%-35%, 10min) to obtained **32** ((7.1 mg, 0.01 mmol, 5.66% yield, 98.57% purity) as a light-yellow solid. $^1$H NMR (400 MHz, DMSO-$d_6$) $\delta$ 9.98 (s, 1H), 8.55 (s, 1H), 8.36 (s, 1H), 7.74 (d, $J$ = 3.6 Hz, 1H), 7.55 (d, $J$ = 8.4 Hz, 1H), 7.31 (dd, $J$ = 1.6, 8.8 Hz, 1H), 6.79 (s, 1H), 6.70 (d, $J$ = 3.2 Hz, 1H), 2.98 (s, 3H), 2.06 (s, 3H). $^{13}$C NMR (101 MHz, DMSO) $\delta$ 168.0, 153.0, 152.6, 151.5, 145.3, 135.2, 134.8, 126.5, 126.1, 120.7, 115.0, 115.0, 105.6, 104.6, 85.5, 77.5, 40.7, 24.1. HPLC Rt = 3.314 min in 8 min chromatography, purity 98.57%.

*N-(1-(3-Cyano-7-(phenylthio)pyrazolo[1,5-a]pyrimidin-5-yl)-1H-indol-6-yl)acetamide (**33**):*

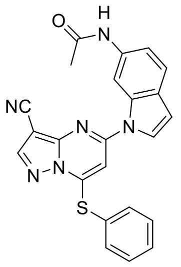

Compound **33** was prepared from **44q** and **46** in the same manner as described for the preparation of **17**. White solid (150.7 mg, 0.35 mmol, 16.97% yield, 99.89% purity). $^1$H NMR (400 MHz, DMSO-$d_6$) $\delta$ 9.99 (s, 1H), 8.85 (s, 1H), 8.53 (s, 1H), 7.89−7.81 (m, 2H), 7.72−7.60 (m, 3H), 7.58−7.48 (m, 2H), 7.24 (dd, $J$ = 1.2, 8.4 Hz, 1H), 6.73 (d, $J$ = 3.2 Hz, 1H), 6.46 (s, 1H), 2.12 (s, 3H). $^{13}$C NMR (101 MHz, DMSO) $\delta$ 168.0, 154.2, 151.5, 149.1, 147.9, 135.9, 135.9, 134.5, 131.5, 130.8, 126.3, 125.8, 124.4, 121.1, 116.0, 113.2, 108.1, 104.9, 98.6, 80.3, 24.0. HPLC Rt = 3.553 min in 8 min chromatography, purity 100%. LCMS $R_t$ = 2.158 min in 4 min chromatography, purity 99.89%, MS ESI *m/z*: 425.3 [M+H]$^+$.
